# Comprehensive Bulk and Single-Cell RNA Sequencing Uncovers Senescence-Associated Biomarkers in Therapeutic Mesenchymal Stem Cells

**DOI:** 10.1101/2025.10.22.683610

**Authors:** Emese Pekker, Erda Qorri, Márton Zs. Enyedi, Valéria Szukacsov, Ferhan Ayaydin, Éva Szabó-Kriston, Bernadett Csányi, Mónika Mórocz, Farkas Sükösd, Endre Kiss- Tóth, Lajos Haracska

## Abstract

**Background:** Mesenchymal stem cells (MSCs) hold great promise in cell therapy, but their effectiveness declines with repeated cell divisions due to senescence. Canines, sharing aging characteristics with humans, serve as a valuable model to study this process in a translational context.

**Methods:** In the present study, we performed an in-depth characterization of senescence in canine MSCs using a combination of morphological, molecular, and transcriptomic analyses. Early (P2) and late-passage (P6) canine MSCs were characterized using a combination of senescence-associated β-galactosidase staining, cell cycle profiling, and both bulk and single-cell RNA sequencing to capture global transcriptional changes.

**Results:** By employing a passage-based *in vitro* approach, the present study demonstrates that late-passage cells (P6) compared to early-passage cells (P2) exhibit hallmark features of senescence, including morphological alterations, elevated SA-β-galactosidase activity, and considerable transcriptional changes. These changes were represented by significant upregulation of established senescence marker genes, alongside potential novel candidates and downregulation of genes associated with cell cycle progression and proliferation. Moreover, single-cell RNA sequencing uncovered heterogeneous distribution of senescent subpopulations, upregulation of SASP-related genes and reduced proliferation markers.

**Conclusions:** Our findings demonstrate that combining classical markers with bulk and single-cell RNA sequencing facilitates senescent cell identification while improving quality control for clinical MSC samples.

## Background

Accumulating evidence suggests that mesenchymal stem cells (MSCs) present a promising option for cell-based therapy. This is attributed to their multipotent differentiation potential, self-renewal capability, extended *ex vivo* proliferation, paracrine potential, and immunoregulatory effects[1, 2]. The characterisation of MSCs’ therapeutic features has revealed significant insights into their regenerative capacity, which has important implications for developing efficient treatments for a wide variety of degenerative conditions[3, 4]. MSCs’ distinctive characteristics make them valuable tools in tissue engineering and regenerative medicine; however, the beneficial functions of MSCs decline with age, a condition which may lead to organ failure and age-related diseases, due to the loss of tissue homeostasis[5, 6].

Despite the presence of MSCs in various tissues, their abundance in the body is relatively limited. Typically, cell therapy protocols require a minimum of 20–100 million MSCs per treatment for autologous transplantation. Consequently, after isolation, *in vitro* expansion of MSCs is required for a period ranging from four to eight weeks before transplantation[7].

Therefore, an adequate number of these cells can be acquired for administration only after expanding them *in vitro* over several population doublings (PDs)[8, 9]. Unfortunately, the proliferative capacity of MSCs decreases during culture expansion[10]. A decreasing division capacity is one sign of’*in vitro* aging’. While embryonic stem cells maintain their proliferative potential *in vitro*[11] and murine MSCs typically do not exhibit signs of *in vitro* senescence[12], human MSCs have shown maximal population doublings of 38-40[13] *in vitro.* At a certain level of PD, MSCs reach the Hayflick limit, cease to proliferate, and enter a senescent state[14].

Therefore, cellular senescence might affect a substantial number of cells during prolonged *in vitro* cultivation. This process is characterized by irreversible growth arrest, which arises from a variety of cellular stresses, such as telomere attrition, DNA damage, oxidative stress, and activation of oncogenes[15]. Senescent cells display alterations in gene expression, including increased expression of cyclin-dependent kinase inhibitors, such as p16 and p21. Senescent cells also exhibit an increase in the lysosomal enzyme senescence-associated-β-galactosidase (SA-β-gal) activity and reduced expression of the nuclear lamina Lamin B1 (LMNB1)[16]. Moreover, senescent cells undergo significant changes in their secretome, releasing pro-inflammatory factors, such as growth factors, cytokines, chemokines, and proteases, collectively known as the senescence-associated secretory phenotype (SASP)[17],[18],[19]. Under normal physiological conditions, SASP factors play a crucial role in tissue repair by orchestrating the recruitment of immune cells and facilitating the removal of damaged tissues[20]. Nevertheless, these secreted factors can induce senescence in the surrounding cells through paracrine signalling, creating an inflammatory microenvironment in their vicinity[21]. MSCs are well-known to exhibit anti-inflammatory properties, whereas senescent MSCs manifest evident pro-inflammatory effects attributed to SASP factors[22]. SASP-related inflammation may contribute to the reduced effectiveness and adverse outcomes of MSC therapies, particularly in elderly patients, driving a debate over their safety and optimal application[23].

Cellular senescence, as an aging mechanism, is a conserved phenomenon across various species[24]. Dogs represent a particularly valuable model for human aging research due to their anatomical and physiological similarities to humans, as well as greater homology in DNA and protein sequences compared to traditional laboratory animals like mice and rats[25]. In addition, canine MSCs (cMSCs) exhibit a somewhat faster rate of PD compared to human MSCs (hMSCs)[26], with a PD time of approximately two days.[27] Research in this area has gained significant interest due to the proinflammatory role of cellular senescence in the clinical application of MSCs, as well as the potential of dogs as an ideal model for identifying novel senescence biomarkers relevant to human aging.

Efforts to characterize senescence biomarkers have identified several markers and techniques. These approaches encompass the assessment of phenotypic changes, cytogenetic techniques, and analyses of genomic and epigenomic profiles[28]. SA-β-gal activity serves as the predominant biomarker for evaluating replicative senescence under *in vitro* conditions, although it is not completely specific[29]. False positive results are caused by confluency, serum starvation, and operator bias[30]. Despite extensive efforts to characterize senescent cells, the identification of reliable and specific senescence markers remains challenging, as existing panels require further optimization to adequately capture the full complexity and heterogeneity of the senescent phenotype[31]. Cellular senescence has been previously described to occur in a tissue-specific manner, and the use of a panel of markers, rather than a single marker, has been proposed for more precise detection of senescent cells. Additionally, high-throughput sequencing methods, such as single-cell/nucleus RNA sequencing, hold great potential for uncovering heterogeneity within senescent cell populations[32].

In this study, we employed a passage-based *in vitro* approach of replicative senescence by culturing cMSCs until replicative exhaustion. With the advancement of molecular techniques, such as bulk RNA sequencing and single-cell RNA sequencing, we sought to investigate the differential gene expression profiles between late-passage cMSCs exhibiting senescent phenotype and early-passage cMSCs. The study aimed to identify potential novel biomarkers and provide a comprehensive characterisation of the heterogeneous nature of replicative senescence at the single-cell level.

## Methods

### MSC expansion and sample collection for the targeted assays

cMSCs were extracted from visceral adipose tissue acquired as surgical waste from healthy dogs that received all routine vaccinations and were regularly examined by a veterinarian. The surgical waste was provided by the veterinarian, with informed consent obtained from dog owners, following standard ovariectomy of the clinically healthy female mixed-breed dogs.

Following the isolation of stromal vascular fraction (SVF) according to the methodology previously described by Kriston-Pál et al.[33], the cells were grown in DMEM/F12 containing 10% foetal bovine serum (FBS) and 50 U/ml penicillin and streptomycin (all provided by Thermo Fisher Scientific, Waltham, Massachusetts, USA). Replicative senescence was induced in the cell culture through serial passaging, involving the progressive subculturing of cells from passage two (P2) through passage six (P6). The cell cultures were grown up to 80% confluence for the six consecutive passages; a medium replacement was performed every three days. Cells were collected from each passage and subsequently stored in liquid nitrogen until processing.

Cell samples underwent assessment for mycoplasma contamination in accordance with the procedures outlined in our prior publication[34]. Proliferating P2-derived cells and P6-derived cells, which underwent replicative senescence, were selected for the study from a 13-month-old donor dog. A total of 10^6^ cells from both P2 and P6 were collected in triplicates for bulk RNA sequencing from the same donor dog. Subsequently, 10^6^ cells derived from P2 and P6 were subjected to single-cell sequencing. In order to determine inter-donor consistency, another dog isolate was also collected and subjected to single-cell sequencing utilizing the methods described above. To evaluate the consistency and generalizability of the observed transcriptomic profiles of early-passage cells, we also included early-passage cells from five additional dog donors.

These supplementary samples were used to determine whether the transcriptional signatures identified in the primary isolate were representative of early-passage cells derived from other individuals. Demographic information of canine isolates used in the study is presented in Supplementary Table 1.

### Tracking morphological changes and calculating population doubling

The morphological changes of the cells were monitored at each passage stage by observing the cells under an EVOS FLoid microscope (Thermo Scientific), and images were captured using transmitted light imaging. PD of each passage was calculated using the formula: log10(N/N_0_)×3.33, where N is the final cell number and N_0_ is the initial number of cells plated. Cumulative PD was determined by summing the PDs of each individual passage.

### SA-β-gal assay

To evaluate senescence of MSCs, SA-β-gal staining was carried out using the SPiDER-βGal kit (Dojindo), according to the manufacturer’s recommendations. A total of 10^5^ cells from P2 and P6 were cultured in triplicates in standard 6-well plates containing coverslips. To minimize the risk of false-positive results caused by confluency, the cells were fixed 24 hours post-culture using 8% paraformaldehyde (diluted in PBS) for 10 minutes at room temperature, followed by three washes with distilled water. For the detection of SA-β-gal, SPiDER-βGal stock solution was diluted 2000 times in McIlvaine buffer (pH 6.0) containing 40 mM citrate-phosphate buffer, 5 mM potassium ferricyanide, 5 mM potassium ferrocyanide, 150 mM sodium chloride, 2 mM magnesium chloride, and distilled water. The cells were incubated in the SPiDER-βGal solution for 40 minutes at 37°C. Nuclei of the cells were stained with DAPI (0.1 mg/ml in PBS) after washing. Coverslips were removed from the plates, drained briefly, and mounted on standard microscope slides using Fluoromount-G antifade mounting medium (Southern Biotech, Birmingham, USA).

### Confocal laser scanning microscopy

Images of SPiDER-βGal-and DAPI-stained cells were acquired with a Leica Stellaris 5 (Leica Microsystems CMS GmbH, Germany) confocal laser scanning microscope using 405 nm laser (DAPI) and 501/525 nm laser lines of white light laser (SPiDER-βGal) with tunable emission filters set to 420–487 nm and 537–614 nm, respectively. Images were taken with an HC PL APO CS2 20× (N.A. 0.75) objective using the Leica Application Suite version X (LAS-X, 4.4.0.24861) software. Identical microscope settings were used for all intensity comparison images. Average SPiDER-βGal intensities were measured using FIJI software (Image J).

Fluorescence intensities were measured using ROI manager, employing a macro tool to draw fixed sized circles according to the code provided by Wayne Rasband on forum.image.sc (https://forum.image.sc/t/changing-point-tool-to-circle-tool/26035/3, accession date: 24.06.2024). Two 30-pixel radius circles were drawn to measure intensities at two different regions of the cytoplasm. For each measurement, a corresponding background intensity was recorded outside the cell, and the actual cellular fluorescence value was normalized to this background. Fluorescence intensity was quantified from exported images (ten randomly selected cell images per replicate) using Fiji (ImageJ) software. Quantitative fluorescence intensity data were collected from 35 cells, with measurements taken from two cytoplasmic regions per cell, yielding a total of 70 measurements per replicate (Supplementary Table 2).

### Assessment of Replicative Potential in BrdU-labelled Cells by Immunofluorescence

Actively growing MSC cells were seeded onto sterile 22 × 22 mm glass coverslips placed in 6-well tissue culture plates at a density sufficient to reach approximately 60-70% confluency the following day. Cells were pulse-labelled with 10 µM 5-bromo-2′-deoxyuridine (BrdU; Sigma-Aldrich) for 24 h to mark actively replicating cells. After labelling, coverslips were transferred (cell side up) to new 6-well plates and washed once with phosphate-buffered saline (PBS). Cells were fixed in freshly prepared methanol:acetic acid (3:1, v/v) for 10 min at room temperature, followed by two washes with PBS. Permeabilization was performed with PBS containing 0.5% Triton X-100 (Sigma-Aldrich) for 10 min. To denature DNA and expose incorporated BrdU, cells were treated with 2.5 N HCl for 1 h at room temperature. Subsequently, cells were washed with 0.5 ml of 0.1 M sodium tetraborate (Na₂B₄O₇, Sigma-Aldrich; pH 8.5) and incubated for an additional 30 min in the same buffer to neutralize residual acid. Coverslips were then washed twice with PBS. Non-specific binding sites were blocked with 5% horse serum and 0.5% Triton X-100 in PBS for 30 min at room temperature. Cells were incubated with mouse anti-BrdU primary antibody (B44 clone; Becton Dickinson, #347580) diluted 1:300 in blocking solution (100 µl per coverslip) and covered with Parafilm to prevent evaporation. The incubation was carried out for 2 h at room temperature. After incubation, coverslips were washed sequentially with PBS and with 1% horse serum/0.2% Tween 20 in PBS. The secondary antibody, Alexa Fluor 488–conjugated goat anti-mouse IgG (Abcam, ab150113), was diluted (diluted 1:400) in blocking solution and applied for 2 h at room temperature. Cells were then washed with PBS and PBS containing 0.2% Tween 20. Finally, coverslips were inverted (cell side down) onto microscope slides using DAPI-containing mounting medium (Fluoromount-G with DAPI; Thermo Fisher Scientific, #00-4959-52). Samples were imaged using using Axioscope Z2 fluorescent microscope (Zeiss, Germany) with a 10x, 20x 40×objective. A minimum of 500 cells from multiple microscopic fields were analysed, and the mean percentage of BrdU-positive cells was plotted.

### Characterisation of cell cycle stages

After incubation in complete growth medium, cells were trypsinized and washed with phosphate-buffered saline (PBS). After washing, cells were resuspended in PBS and centrifuged at 500 rcf for 10 min. After the first centrifugation step, 100% ethanol was added dropwise to the pellet, which was then resuspended in PBS. After the second centrifugation step at 500 rcf for 10 min, the cell pellet was resuspended in RNase solution (100 µg/ml) and PBS. Propidium iodide (1 mg/ml) was added to the cell suspension which was then analysed using the CytoFLEX S flow cytometer (Beckman Coulter). A total of 10000 cells were gated based on forward scatter area (FSC-A) versus side scatter area (SSC-A) dot plots. Singlets were identified and gated using PE versus PE-A dot plots.

### RNA isolation for bulk RNA sequencing and quantification

RNA was extracted from three parallel samples obtained from passages 2-and 6-derived cells (10^6^ cells each) using the RNeasy kit (Qiagen, Hilden, Germany), following the manufacturer’s recommendations. Quantification of extracted RNA was performed using the Qubit® 2.0 Fluorometer (Invitrogen, Milan, Italy) and the Qubit® RNA HS Assay Kits (Life Technologies, Carlsbad, CA). Total RNA samples were quality checked with a BioAnalyzer 2100 instrument using the Agilent RNA 6000 Nano Kit (Agilent Technologies USA, Cat. No. 5067-1511). The RNA integrity number (RIN) values obtained were as follows: passage 2 - replicate 1: 8.60, replicate 2: 9.50, replicate 3: 9.60; passage 6 - replicate 1: 8.40, replicate 2: 9.30, replicate 3: 8.10.

### Library construction and quality control for bulk RNA sequencing

Next-generation sequencing (NGS) library preparation was carried out on 500 ng RNA for each sample, using the NEBNext Ultra™ II Directional RNA Library Prep Kit for Illumina (NEB #E7760) with NEBNext Poly(A) mRNA Magnetic Isolation Module (NEB #E7490).

Sequencing-ready libraries were quality control checked with a BioAnalyzer2100 instrument using High Sensitivity DNA Chip (Agilent Technologies USA, Cat. No. 5067-4626). NGS was carried out on the NextSeq 500 sequencing system with NextSeq 500/550 Mid Output Kit v2.5 (150 Cycles) chemistry (Illumina, Inc. USA, Cat. No. 20024904) targeting 20M reads/sample.

### Bulk RNA sequencing data processing

The quality of the raw data was assessed using FastQC (v0.11.9)[35]. Adapter sequences and low-quality reads were removed using FASTP (v0.12.4)[36]. The cleaned paired-end reads were pseudo-aligned to the *Canis lupus familiaris* reference transcriptome (CanFam3.1 build, Ensembl release 99) using *kallisto quant* (v0.46.1) with default parameters[37]. Quality control of the raw data and pseudoalignment metrics can be found in the Supplementary Table 3A. The reference transcriptome index was generated with the *kallisto index* command. Transcript abundance estimates were summarized to gene level using the tximport package[38], filtered with the filterByExpr function (edgeR)[39], and normalized using the trimmed mean of M-values (TMM) method in edgeR[39]. Differential gene expression analysis was performed using limma-voom[40], and genes with a log2 fold change (log2FC) > 1 and adjusted p-value < 0.01 were considered significantly differentially expressed. These genes were used as input for the Gene Ontology analysis.

Gene overrepresentation analysis was performed using the enrichGO function from the clusterProfiler package[41]. Fold enrichment values for each GO term were calculated according to the formula provided in the clusterProfiler article[41] and then ranked from highest to lowest. For the KEGG enrichment analysis, the *search_kegg_organism*function was first used to retrieve the KEGG abbreviation for *Canis lupus familiaris*, and the *enrichKEGG* function was then utilized to identify enriched pathways in the dataset, with significant pathways determined by a p-value cutoff of 0.05.

To assess whether the transcriptome profiles of our early-passage-derived (P2) samples are similar to other early-passage cells from five different dog donors, we performed a Transcripts Per Million (TPM)-based correlation analysis. The reads from passage 3-derived samples from five different donors were pseudo-aligned to the reference *Canis lupus familiaris* transcriptome (Ensembl release 99) using Kallisto with default settings. The resulting TPM values were then compared to those from passage 2 and its technical replicates.

### Preparation of single-cell suspensions for single-cell sequencing

Cryovials containing 10^6^ cells were removed from liquid nitrogen storage and immediately thawed in water bath at 37°C for 1 min. Subsequently, cell suspensions were transferred to 10 ml of warm complete medium (DMEM/F12 containing 10% foetal bovine serum (FBS) and 50 U/ml penicillin and streptomycin) and centrifuged at 300 rcf for 5 min. After discarding the cell supernatant, cell pellet was resuspended in 10 ml of complete medium. Following another centrifugation and removal of the supernatant, 1 ml of 1x PBS + 0.04% BSA was gently added to the cell pellet and resuspended. After centrifugation, a cell concentration of 700 cells/µl was achieved by resuspending the pellet in 1x PBS + 0.04% BSA. Cell concentration and viability were assessed following the initial and third centrifugation steps using Bürker chamber by diluting the cells with trypan blue. After adjusting the cell concentration of the single-cell suspension to 700 cells/µl, it was loaded into the Chromium Controller system (10x Genomics, Pleasanton, CA, USA), targeting 10000 cells/sample. mRNA capture, cDNA amplification, and library construction were conducted according to the manufacturer’s instructions using Single Cell 3ʹ GEM, Library & Gel Bead Kit v3.1 (10x Genomics, 1000269). The generated mRNA libraries were sequenced using the Illumina NovaSeqX Plus system, targeting 20000 read pairs per cell.

### Single-cell read mapping and quantification

Raw single-cell data was processed using the Cell Ranger (v.7.2.0) software by 10x Genomics[42]. Reads were aligned to the *Canis lupus familiaris* Cfam3.1 genome (updated on 2019-11-19), retrieved from the Ensembl database[43]. Quality metrics of this process are accessible in Supplementary Table 3B. The filtered feature-barcode matrices were processed in R (version 4.4.1)[44] using the Seurat package (version 4.4.0)[45]. To ensure high quality data for downstream analysis, only those cells with ≥550 detected genes, ≥650 unique molecular identifiers (UMIs), and less than 10% mitochondrial contamination were retained. We then performed a gene-level filtering to reduce noise from lowly expressed genes. Specifically, we excluded genes with zero counts, and among the remaining ones, retained only those expressed in ≥10 cells for downstream analyses.

After filtering, data normalization was performed using the *NormalizeData* function with default settings. To identify the principal components (PCs) contributing most to variance, the Seurat object was scaled with the *ScaleData* function, followed by principal component analysis (PCA). The data were then clustered using the *FindNeighbours* and *FindClusters* functions, and the clusters were visualized using the Uniform Manifold Approximation and Projection (UMAP) technique. Multiple resolutions (0.2, 0.4, 0.6, 0.8, 1.0, and 1.4) were tested to determine the optimal number of subclusters, and a resolution of 0.2 was selected following a close visual inspection and consideration of the study’s objectives.

To identify genes expressed in each cluster from the P2 and P6 samples, the *FindAllMarkers* function was used with thresholds of Log fold change (logFC) > 1 and adjusted p-value < 0.05 for marker gene selection.

Additionally, to ensure reproducibility and allow for cross-donor comparison of the identified markers, a second single-cell RNA-sequencing run was performed on a second dog donor and analysed independently, using the pipeline described above. After data quality assessment, only cells with ≥650 UMIs and ≥550 detected genes were retained for further analysis (Supplementary Table 4). Clustering was performed on the first ten principal components, and UMAP was applied to visualize the separation of cells according to the origin of the sample.

Following this analysis, both runs were merged into a single Seurat object and integrated to correct for technical variation using Seurat’s canonical correlation analysis (CCA) framework with SCTransform normalization (3000 variable features). The merged object was split by donor and passage identity: P2-R1 and P6-R1 (Donor1/Run1) and P2-R2 and P6-R2 (Donor2/Run2).

Final integration was performed using 30 principal components, which provided optimal alignment between equivalent passages, while preserving clear biological differences between early and late passages. Cells were subclustered at a resolution of 0.2, and the contribution of each passage to the resulting clusters was visualized using barplots. Mitochondrial transcripts were excluded during the preprocessing of the second run; to ensure consistency, mitochondrial genes were also excluded from the filtered object of the first run prior to integration.

### Analysis of publicly available datasets

To investigate the expression patterns of the identified senescence markers across different organisms, the datasets from Casella et al. (2019)[46] (GEO ID: GSE130727) and Wang et al. (2022)[47] (GEO ID: GSE179880) from human and mice samples, respectively, were analysed. The analysis was carried out as previously described in the section “Bulk RNA-sequencing data analysis”, with significance thresholds set according to the criteria reported in the respective publications.

An additional single-cell RNA-sequencing dataset from Fard et al.[48] (2023: GEO accession GSE200157) was included in the analysis. From the available samples, only the T0 and T2 time points were retrieved and compared. Droplets were initially filtered using DropletUtils[49] and quality control was performed as previously described applying the following thresholds: UMIs ≥ 600, features ≥ 500 and ≤ 6000, and mitochondrial content < 20%. The filtered data were normalized using SCTransform and integrated with Harmony[50] based on 30 principal components. The expression levels of the selected markers were subsequently examined using dot plots.

### Statistical Analyses and Software

The differences in β-gal activity between P2 and P6 were assessed using the Wilcoxon Rank Sum test. Prior to analysis, data distribution was visually inspected and formally tested for normality using the Shapiro-Wilk test. As the data do not follow a normal distribution, Wilcoxon Rank Sum test was selected. All statistical analyses including the Spearman rank correlation analysis of the TPM expression profiles were performed in R (version 4.4.1). Additionally, visualizations, including PCA, donut charts, and volcano plots were generated using the ggplot2[51] and EnhancedVolcano packages[52]. Heatmaps were generated using the pheatmap package unless otherwise specified. Workflows and figure panels were assembled using BioRender.

For the BrdU experiments, statistical differences between P2 and P6 were evaluated using an unpaired t-test. Prior to analysis, data normality was assessed using the Shapiro–Wilk test, which confirmed that the samples did not significantly deviate from a normal distribution. Therefore, an unpaired t-test was applied to determine differences between the samples. The mean values obtained from microscopic field counts were plotted with standard deviations.

### Cell cycle analysis

To determine the cell cycle stage of the cells in both samples, the *CellCycleScoring* function from Seurat was utilized[45]. The function calculates enrichment scores for the S and G2M phases, assigning each cell to one of three phases: G2M, S, or G1. Cells with negative scores for both the G2M and S phases are classified as G1 cells[53]. The list of human cell cycle marker genes compiled by Tirosh, I. et al. was retrieved from Seurat[54]. For *Canis lupus familiaris*, the cell cycle gene list was generated through ortholog searches using the guidelines provided in the tutorial https://github.com/hbc/tinyatlas. Based on the cell cycle classification, the percentage of cells in each phase (G1, S, and G2M) was calculated.

### Co-occurrence analysis

To identify novel senescence-related markers, we devised a co-occurrence approach based on the hypothesis that a cell must express key senescence markers (cited in over 20 scientific publications on cellular senescence) such as CCND1, IGFBP2, CDKN1A, TGFB2, along with at least one of the top ten genes upregulated in P6 (CDKN2A, IGFBP7, CRYAB, ITGA2, PTN, NDUFA4L2, COL11A1) identified through the *FindAllMarkers* function. After identifying these cells, we calculated the percentage of senescent cells within each subcluster.

## Results

### cMSCs undergoing replicative senescence display altered cell morphology and increased β-galactosidase activity

Proliferating MSCs from P2 were cultured until P6, where they reached replicative exhaustion, leading to cellular senescence. P2 cells exhibited a population doubling level (PDL) of 2.7 over a 7-day culture period, whereas P6 cells showed a PDL of 0.75 over a 14-day culture period (Supplementary Figure 1), reaching a cumulative PDL of 8.67. Proliferating cells derived from P2 and P6-derived cells undergoing replicative exhaustion were selected for the present study.

We observed that late-passage cells derived from P6 exhibited enlarged and flattened morphology, characteristic of senescent cells, in contrast to the P2-derived proliferating cells (Supplementary Figure 2). Cellular senescence was confirmed by investigating the senescent phenotype of the P6-derived cells; P2 and P6 cells were treated with Spider β-Gal. In P6-derived cells, an increase in fluorescence signal was observed, indicating replicative senescence.

Representative images can be seen in Figure 1A. A significantly increased β-galactosidase activity was observed in P6 cells compared to P2 cells, increasing from 6.8 in P2 to 40.3 in P6 (Figure 1B).

**Figure 1.**
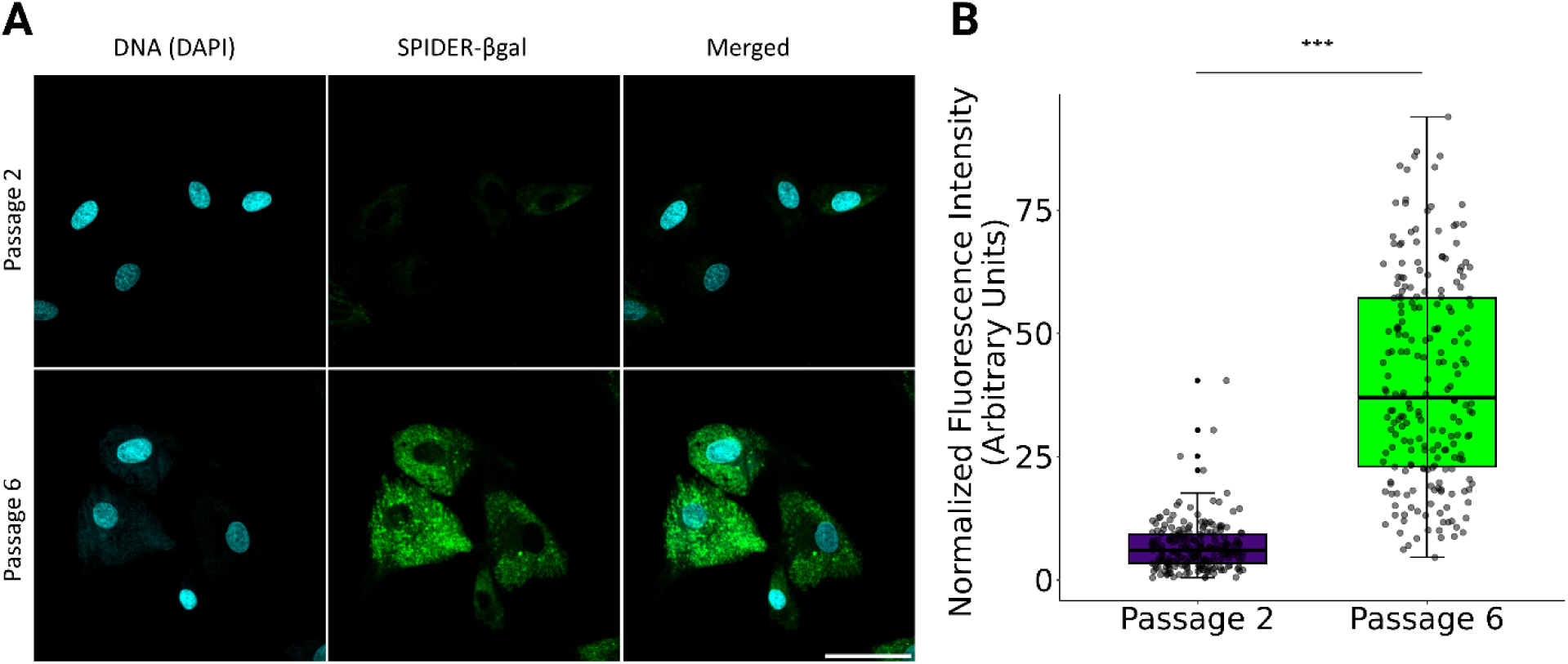
SA-β-gal assay. A. Representative images of SA-β-gal staining assay. Scale bar is 50 µm **B.** Quantitative fluorescence intensity data were obtained from 35 cells, measuring two cytoplasmic regions per cell, resulting in a total of 70 measurements per replicate. For each measurement, background fluorescence outside the cell was recorded and used for normalization. The boxplot whiskers extend from the third and first quartiles to the largest and smallest values, respectively, within 1.5 x the interquartile range (IQR), where IQR represents the range between the first and third quartiles.

The observed difference in fluorescence intensity was considered statistically significant, with a p-value <2.2e-16.

To further validate these observations, we next assessed proliferative capacity using the BrdU incorporation assay. Analyses were performed at passages 2 and 6, once cultures reached 50– 60% confluence to ensure comparable growth conditions. Consistent with the β-galactosidase results, BrdU incorporation revealed a pronounced reduction in proliferation with increasing passage number: 82% of P2 cells incorporated BrdU, compared to only 25% of P6 cells (Figures 2A and 2B).

**Figure 2.**
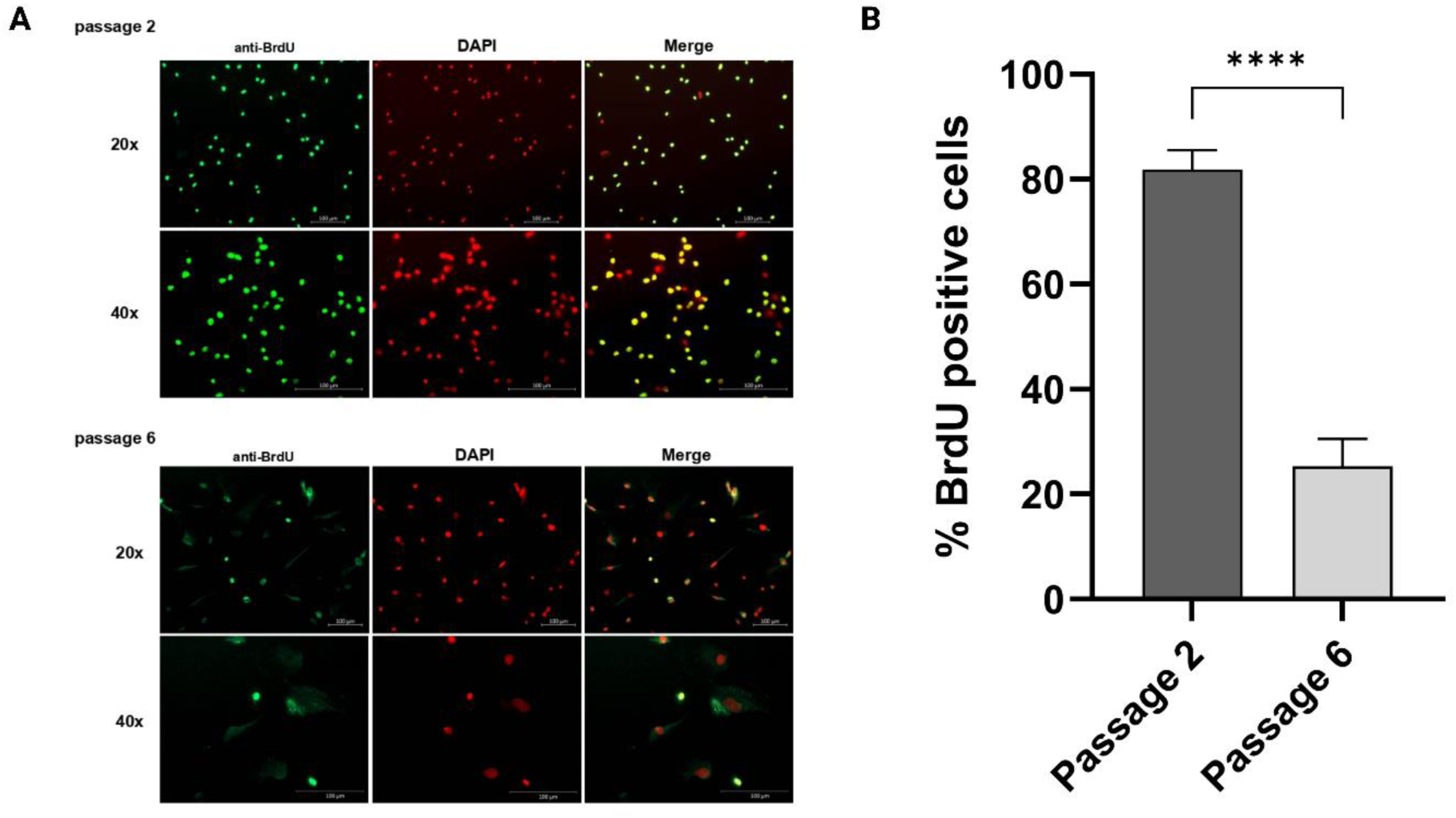
Reduced proliferative activity in late-passage cells, determined by BrdU incorporation. **A.** Representative immunofluorescence images of cells at passage 2 (top) and passage 6 (bottom) stained with anti-BrdU antibody (green) and DAPI (red) to visualize the nuclei. Merged images show BrdU-positive nuclei in yellow. Images were acquired at 20× and 40× magnifications. Scale bars: 100 μm. **B.** Quantification of BrdU-positive cells expressed as a percentage. Data represents the mean ± standard deviation from microscopically counted fields.; ****p < 0.0001.

2A-B). This sharp decline in DNA synthesis activity confirms that late-passage MSCs undergo significant proliferative arrest, which might be in line with the onset of replicative senescence.

### Comprehensive profiling of differentially expressed genes (DEGs), associated processes, and pathways in MSC senescence

To evaluate replicative senescence-mediated gene expression changes in MSCs, we initially carried out bulk RNA sequencing. MSCs were harvested across six consecutive cell passages, with cells from passages P2 and P6 chosen for detailed analysis. Bulk RNA sequencing was performed on P2-and P6-derived cells in triplicates, following RNA extraction and library preparation, using the Illumina NextSeq 500 platform. The sequencing data were subsequently subjected to comprehensive bioinformatic analysis (Figure 3A). The gene expression profiles of MSCs derived from P6 were compared to those of P2-derived cells using three independent experimental replicates. Principal Component Analysis (PCA) revealed a clear separation between the early-and late-passage samples (Supplementary Figures 3A and 3B). A total of 1091 genes were identified as DEGs, including 533 upregulated genes (Supplementary Table 5) and 558 downregulated genes (Supplementary Table 6 and Figure 3B).

**Figure 3.**
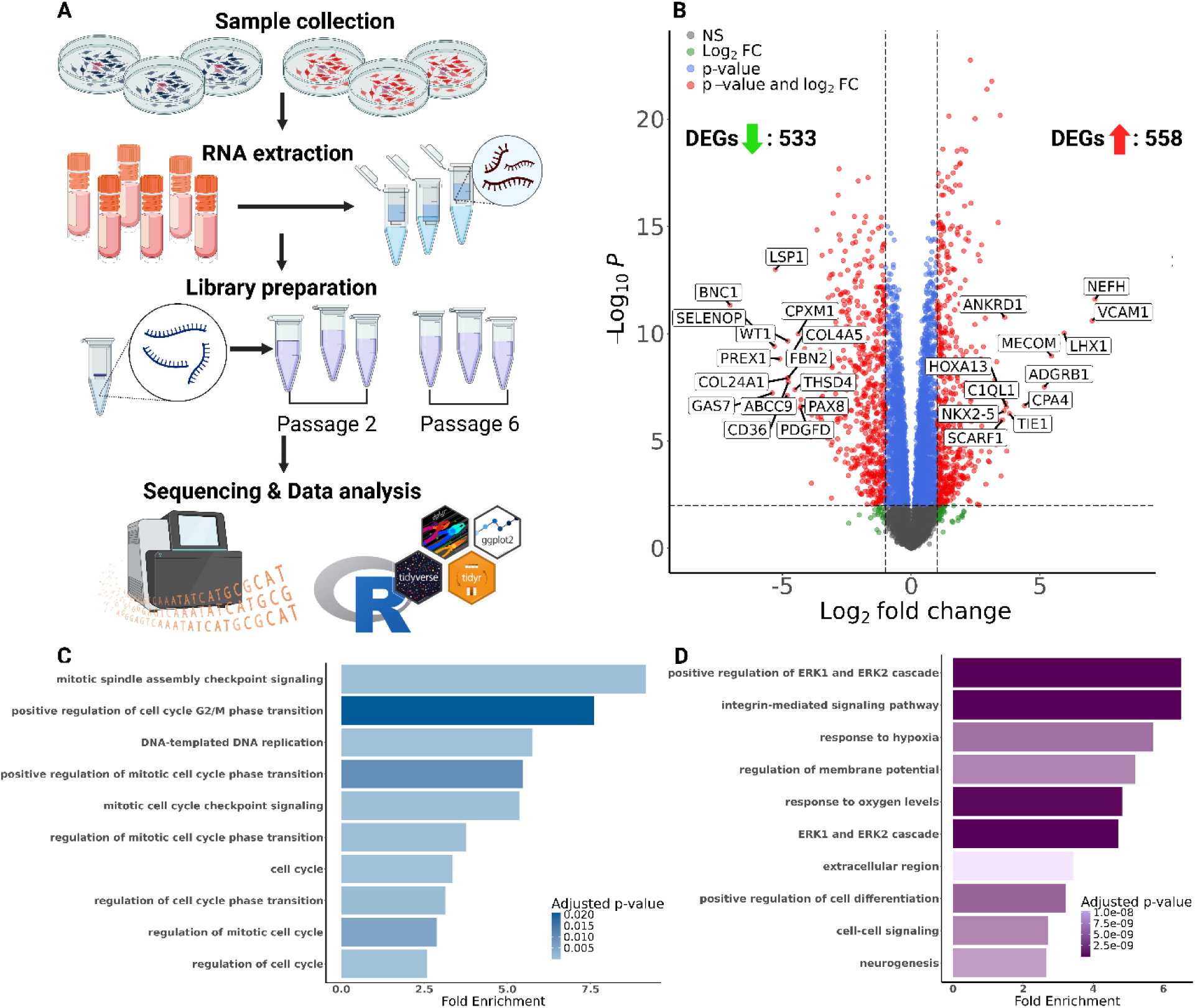
Bulk RNA sequencing-based transcriptomic analysis. A. Schematic representation of the workflow for bulk RNA seq-based assay. MSCs were collected from six consecutive cell passages. P2-and P6-derived cells were selected for the study. Bulk RNA sequencing of P2-and P6-derived cells was carried out in triplicates, following RNA extraction and library preparation using Illumina NextSeq 500, and the results were analysed bioinformatically. Created in BioRender. Kiss, E. (2025) https://BioRender.com/2i3n4gv. **B.** Volcano plot highlighting the top identified DEGs in P6-derived samples, compared to P2 samples. Red data points indicate those genes that were significantly upregulated (right) or downregulated (left). X-axis=Log_2_FC, horizontal dashed line indicates cutoff for P-value<0.01, while the vertical dashed lines indicate the cutoff for fold change of 2. **C.** Results of GO enrichment analysis for the downregulated differentially expressed genes. X-axis = Fold enrichment values. Y-axis=relevant enriched GO terms for different biological processes; colour intensity of bars based on p-value. **D.** Results of GO analysis for the upregulated DEGs. X-axis = Fold enrichment values. Y-axis=relevant enriched GO terms for different biological processes; colour intensity of bars based on p-value.

Among the upregulated DEGs, the most prominent were NEFH, VCAM1, LHX1, MECOM, ADGRB1, CPA4, TIE1, HOXA13, C1QL1, ANKRD1, and NKX2-5, along with two unannotated transcripts (ENSCAFG00000044398 and ENSCAFG00000031632). Among these, the NEFH (neurofilament heavy chain) gene exhibited the highest logFC, with a value of 7.12. CCND1 (Cyclin D1), CDKN1A (Cyclin Dependent Kinase Inhibitor 1A), and CDKN2A (Cyclin Dependent Kinase Inhibitor 2A) also showed significant upregulation in P6, reflecting cell cycle arrest. In contrast, BNC1 (Basonuclin Zinc Finger Protein 1) exhibited the most pronounced decrease in expression levels (logFC =-7), along with GAS7, SELENOP, LSP1, PREX1, ABCC9, COL24A1, WT1, COL4A5, CD36, FBN2, THSD4, CPXM1, PDGFD, and PAX8, indicating diminished metabolic and extracellular matrix activity, consistent with loss of proliferative capacity. The top differentially expressed genes based on log2FC values are illustrated in Figure 3B, with downregulated DEGs shown on the left and upregulated DEGs on the right.

To determine the similarity between the transcriptome profiles of our early-passage-derived (P2) samples and early-passage cells from five different dog donors, we conducted a correlation analysis based on TPM values. Spearman rank correlation revealed that the P2 samples show strong correlations (≥0.96) with P3 samples from five different dog donors (Supplementary Figure 4), suggesting that the transcriptome profiles of early-passage cells are highly similar to those of passage 3-derived cells. Overall, these results indicate that the transcriptomic profile of our early-passage-derived (P2) samples is comparable to early-passage cells from other dog donors, supporting their reliability and consistency for further analysis.

To assess the functional significance of the DEGs, we performed GO and KEGG enrichment analyses separately for up-and downregulated genes. Downregulated genes showed significant enrichment in 142 GO terms (Supplementary Table 7), primarily related to cell cycle regulation and DNA replication, consistent with reduced proliferative capacity (Figure 3C). In contrast, the 178 enriched GO terms for upregulated genes (Supplementary Table 8) were linked to ERK signalling, hypoxia response, cell differentiation, and neurogenesis, reflecting response to replicative stress of these cells (Figure 3D).

To further examine the functions of the identified differentially expressed genes within specific pathways, we conducted KEGG pathway analysis. A total of 23 pathways exhibited significant enrichment, among which 15 were linked to the upregulated DEGs (Supplementary Table 9).

Notably, the most significantly enriched pathways included hypertrophic cardiomyopathy, dilated cardiomyopathy, and hematopoietic cell lineage. Additionally, the HIF-1 signalling pathway, the PI3K-Akt signalling pathway, and pathways related to cancer also demonstrated significant enrichment. The downregulated DEGs were assigned to eight pathways (Supplementary Table 10); among them, DNA replication, cell cycle and progesterone-mediated oocyte maturation showed the highest representation.

### UMAP analysis reveals distinct clustering of early and late passage cMSCs

To investigate the molecular mechanisms underlying the heterogeneity in MSC senescence, we performed a comprehensive single-cell transcriptomic analysis on cMSCs subjected to replicative senescence. To characterize the differences between senescent cells from P6 and actively dividing cells from P2, as well as the heterogeneity within the senescent cell population, we conducted single-cell gene expression profiling using the specific 10x Single Cell 3ʹ GEM pipeline, and subsequently caried out sequencing using the Illumina NovaSeqX Plus system (Figure 4A).

**Figure 4.**
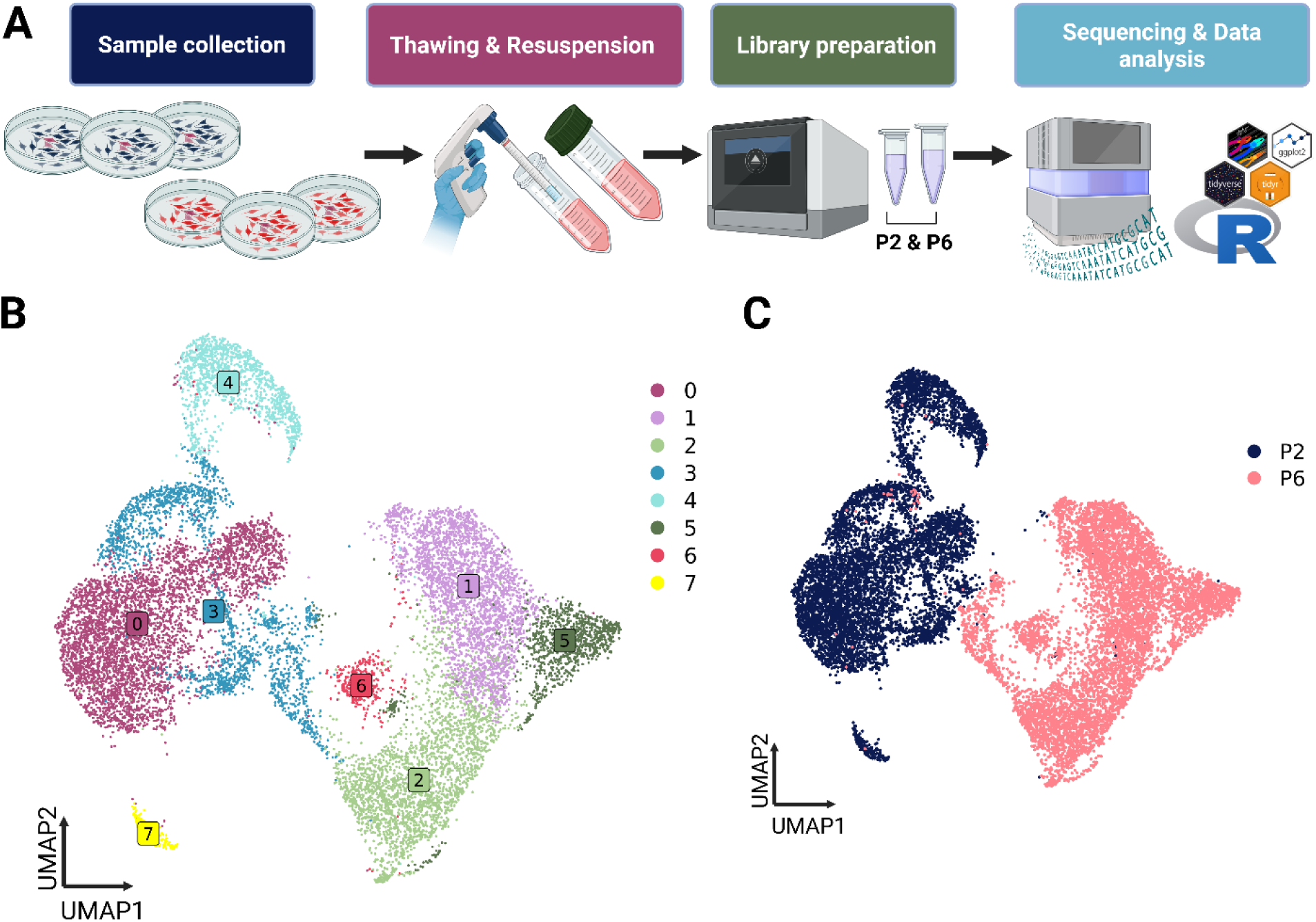
Single-cell RNA sequencing workflow and UMAP clustering analysis of cells derived from P2 and P6. A. MSCs were collected from six consecutive cell passages, and P2-and P6-derived cells were selected. Single-cell RNA sequencing was carried out using Single Cell 3ʹ GEM, Library & Gel Bead Kit v3.1 and sequenced using the Illumina NovaSeqX Plus system, and the results analysed bioinformatically. Created in BioRender. Kiss, E. (2025) https://BioRender.com/lncnz90. **B.** Distribution of the identified clusters. **C.** Distribution of the clusters based on the origin of the sample (P2= Passage 2, P6= Passage 6).

UMAP analysis of the dataset comprising cells derived from P2 and P6 unveiled the presence of eight distinct clusters (see Figure 4B) organized in two major cell populations. Subsequent plotting of cells according to the origin of the sample demonstrated that the two major cell populations are represented by a distinct separation between early-passage cells and late-passage senescent cells (Figure 4C). The early-passage cell population (P2) predominantly comprised clusters 0, 3, 4, and 7, while the late-passage cell population (P6) was dominated by clusters 1, 2, 5, and 6. Cells in clusters 0, 3, 4, and 7 constituted 99.7%, 77.5%, 99.2%, and 100%, respectively, of the P2-derived cell population. Cells in clusters 1, 2, 5, and 6 comprised 99.3%, 99.6%, 99.3%, and 99.6%, respectively, of the cells derived from P6, suggesting considerable differences in the transcriptional profile of early-and late-passage cell populations. Details are shown in Supplementary Figure 5.

The housekeeping genes Actin Beta (ACTB), Beta-2-Microglobulin (B2M), Glyceraldehyde-3-Phosphate Dehydrogenase (GAPDH), and Phosphoglycerate Kinase 1 (PGK1) were analysed across the clusters, and all were found to be expressed throughout (Supplementary Figure 6). The number of cells expressing the abovementioned genes per cluster can be found in Supplementary Table 11. Expression of established MSC markers – CD44, THY1 (CD90), FN1, ALCAM, ITGB1 (CD29), and ENG (CD105) – was detected across the clusters, supporting the origin and identity of the analysed MSC cell populations (Supplementary Figure 7).

### Cell cycle stages and proliferative properties of actively dividing and senescent cMSCs at the single-cell level

One of the main senescence-associated features includes cell cycle arrest of cells in the G1 phase. To evaluate cell cycle arrest during the emergence of replicative senescence, flow cytometry was performed. P2-and P6-derived cells were collected and stained with propidium iodide. The cells were subsequently analysed using flow cytometry. As shown in Figure 5A, P6 cells exhibited a marked increase in G1 phase accumulation (83.2%) compared to P2 cells (71.5%).

**Figure 5.**
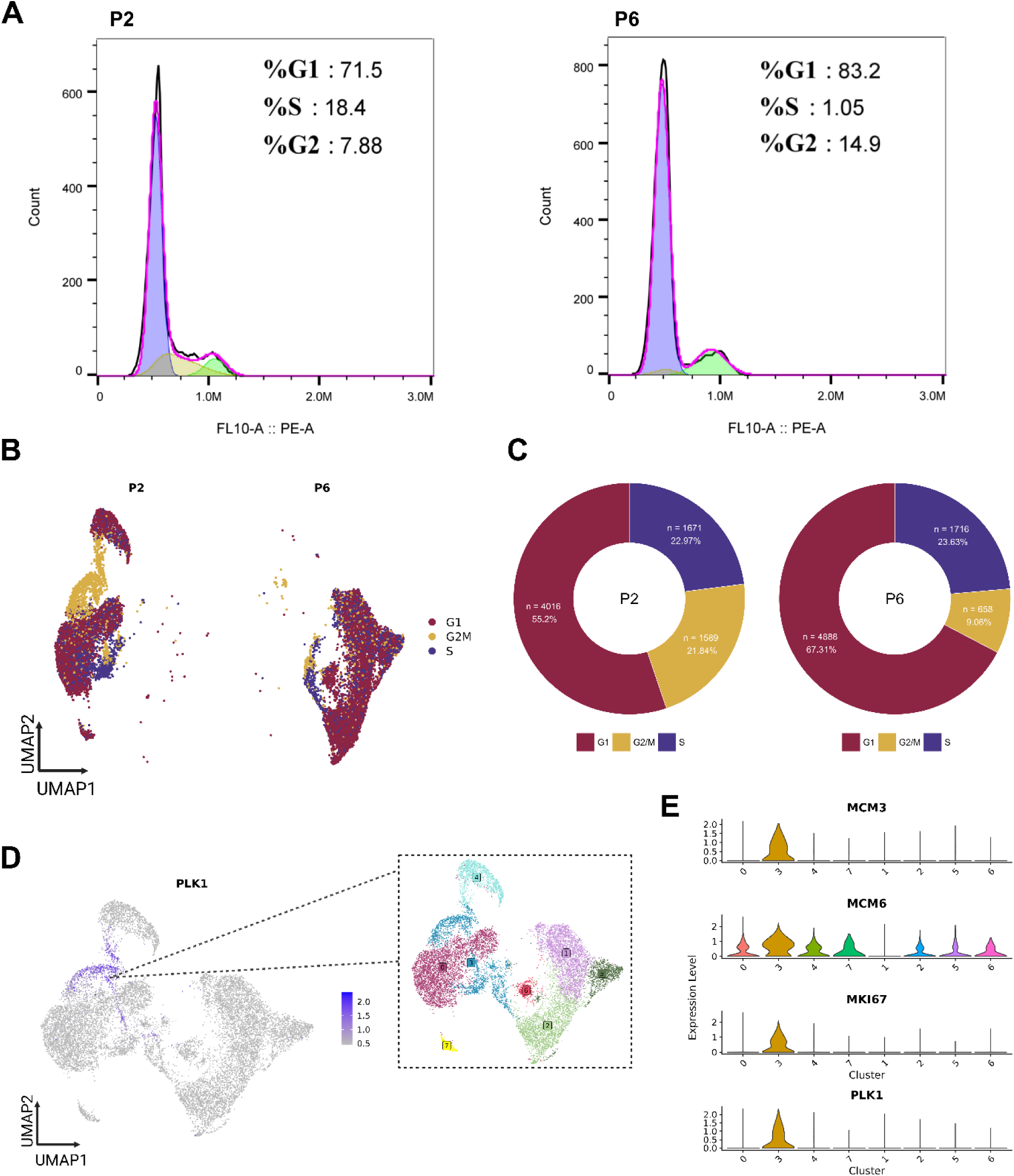
Cell cycle stages and proliferation dynamics within the clusters. **A.** The cell cycle of the sorted cells was assessed by flow cytometry using propidium iodide staining. Representative images of P2 (left) and P6 (right) cells are shown, highlighting that P6 cells are represented by a considerably higher proportion of cells in the G1 phase. **B-C**. Cell cycle stages of the cell population and percentages are displayed separately for P2 (left) and P6 (right), as determined by Cell Scoring analysis using the Seurat package. **D**. Feature plot representing the expression of proliferation marker PLK1 among the clusters. Inset on the right highlights cluster distribution **E.** Expression of transcripts for cell proliferation markers and division regulators PLK1, MKI67, MCM3, and MCM6. The clusters are arranged based on the origin of the sample (P2 followed by P6).

We further employed the *CellCycleScoring* function in the Seurat package to conduct cell cycle analysis and identify the distribution of G1-arrested senescent cells across clusters (Figure 5B), using transcriptomic signatures specific to distinct cell cycle phases. In line with flow cytometry results, cell scoring estimated that 67.31% of cells derived from P6 were in the G1 phase compared to 55.2% of cells from P2 (Figure 5C). Consistent with this, the specific late-passage clusters (clusters 1, 2, 5, and 6) demonstrated diminished expression of transcripts for cell proliferation markers and division regulators, including PLK1 (Figure 5D), MKI67, and MCM3, and displayed decreased expression for MCM6, further supporting that subsets within these cell populations were likely not undergoing active division. Notably, PLK1, MKI67, and MCM3 were expressed exclusively in cluster 3 from the early-passage cell population, while MCM6 exhibited elevated expression relative to clusters 0, 4, and 7 within the early-passage cells, further underscoring that cluster 3 is characterized by a high proportion of proliferating cells (Figure 5E). These results strongly indicate that P6-derived cells are predominantly G1-arrested, whereas early-passage P2 cells exhibit elevated expression of the proliferation marker MCM6.

Additionally, early-passage P2 cells include a specific cluster with pronounced expression of cell proliferation markers and division regulators, supporting the active proliferative status of these cells from cluster 3.

### Evaluation of senescence-related pathways in cMSCs

We evaluated the expression of genes associated with the enriched GO terms identified through bulk RNA sequencing, focusing on three key pathways highly relevant to cellular senescence: the ERK1 and ERK2 cascade, the integrin-mediated signalling pathway, and response to oxygen levels. Consistent with the observations from bulk RNA sequencing, the late-passage cell populations represented by clusters 1, 2, 5, and 6 showed elevated expression of the ERK1 and ERK2 cascade-related genes APP, EDN1, and CCL5. Notably, cluster 6 showed reduced expression of EDN1 and CCL5 compared to other late-passage clusters (Figure 6A). Another important significantly enriched GO term in P6 samples was the integrin-mediated signalling pathway. Single-cell analysis revealed elevated expression of the pathway-associated genes TIMP1, ITGA1, and ITGA2 in specific late-passage clusters. Interestingly, ITGA1 exhibited higher expression in cluster 7, while ITGA2 was more prominently expressed in clusters 2 and 6 (Figure 6B). Elevated oxygen concentrations frequently trigger the onset of senescence, underscoring the significance of oxygen level responses in cellular senescence[55]. The expression profiles of genes associated with the oxygen-response pathway demonstrated elevated levels in clusters derived from P6, notably including TGFB2. However, EDN1 exhibited downregulation in cluster 6. Additionally, CRYAB was most highly expressed in cluster 7 (Figure 6C).

**Figure 6.**
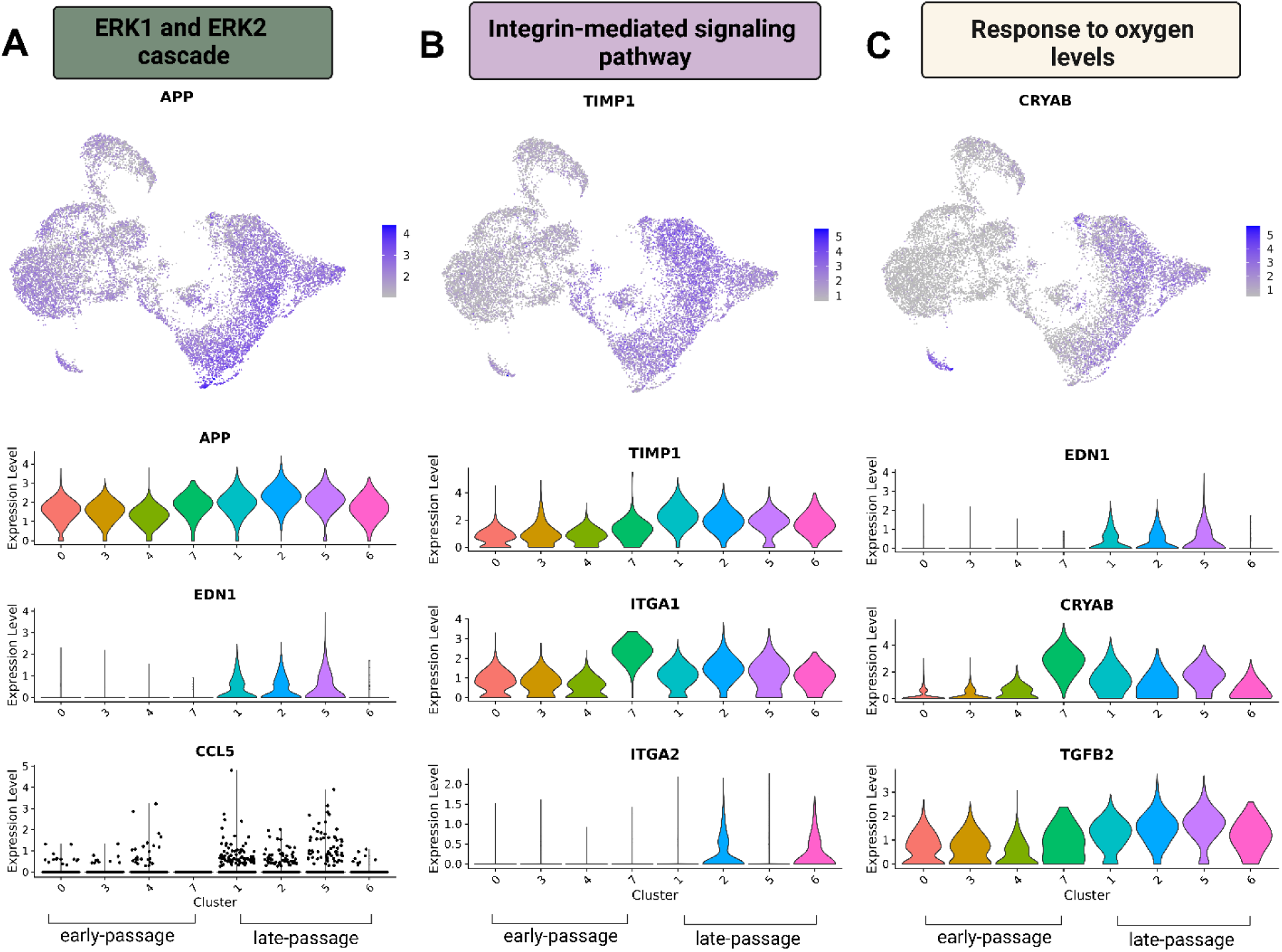
**Evaluation of the significantly enriched GO terms, emphasizing three critical pathways associated with cellular senescence at the single-cell level**. **A.** the ERK1 and ERK2 cascade: feature plot illustrating the expression of the APP gene (top), violin plots showing the expression levels of the ERK1 and ERK2 cascade-related genes APP, EDN1, and CCL5 among the clusters (bottom). **B**. Integrin-mediated signalling pathway: feature plot illustrating the expression of the TIMP1 gene (top), violin plots showing the expression levels of the integrin-mediated signalling pathway-related genes TIMP1, ITGA1, and ITGA2 (bottom). **C.** Response to oxygen levels: feature plot illustrating the expression of the CRYAB gene (top), violin plots showing the expression levels of response to oxygen levels GO term-related genes EDN1, CRYAB and TGFB2 (bottom). The clusters are arranged based on the origin of the sample (P2 followed by P6).

### Characterisation of the unique transcriptional profiles of clusters and identification of shared senescence-associated and SASP-related signatures

To characterize the transcriptional profiles of the clusters, we employed the *‘FindAllMarkers’* function to identify genes with the highest representation uniquely associated with each cluster (details are shown in Supplementary Table 12). Remarkably, cluster 3 exhibited a distinct transcriptional signature, characterized by the top five highly expressed genes that showed diminished expression in the other clusters. These included CDC20, a key regulator of the metaphase-to-anaphase transition during cell division, DIAPH3, which influences cell motility and adhesion, STMN1, which modulates signals of the cellular environment, and RRM2 and HMGB2, involved in cell proliferation, demonstrating that cells from cluster 3 are engaged in active cell division and proliferation. Late-passage cell clusters 1, 2, 5, and 6 exhibited transcriptomes associated with cellular senescence. These clusters demonstrated elevated expression of several components of SASP, including IGFBP2, IGFBP7, and PAPPA, as well as the senescence-related genes CCND1 and CRYAB. However, the expression of IGFBP2 was reduced in cells from cluster 6, whereas IGFBP7 displayed prominent expression in cluster 1 derived from P6, with notably high levels of these SASP factors also detected in cluster 7.

CCND1 exhibited strong expression in P6 clusters 1, 2, 5, and 6, while CRYAB showed prominent expression in clusters 1 and 5, with particularly high levels observed in cluster 7. Interestingly, OGN (Osteoglycin), described as a negative regulator of cellular senescence[56], exhibited elevated expression across all late-passage clusters (1, 2, 5, and 6). The top five identified marker genes are represented in Figure 7A.

**Figure 7.**
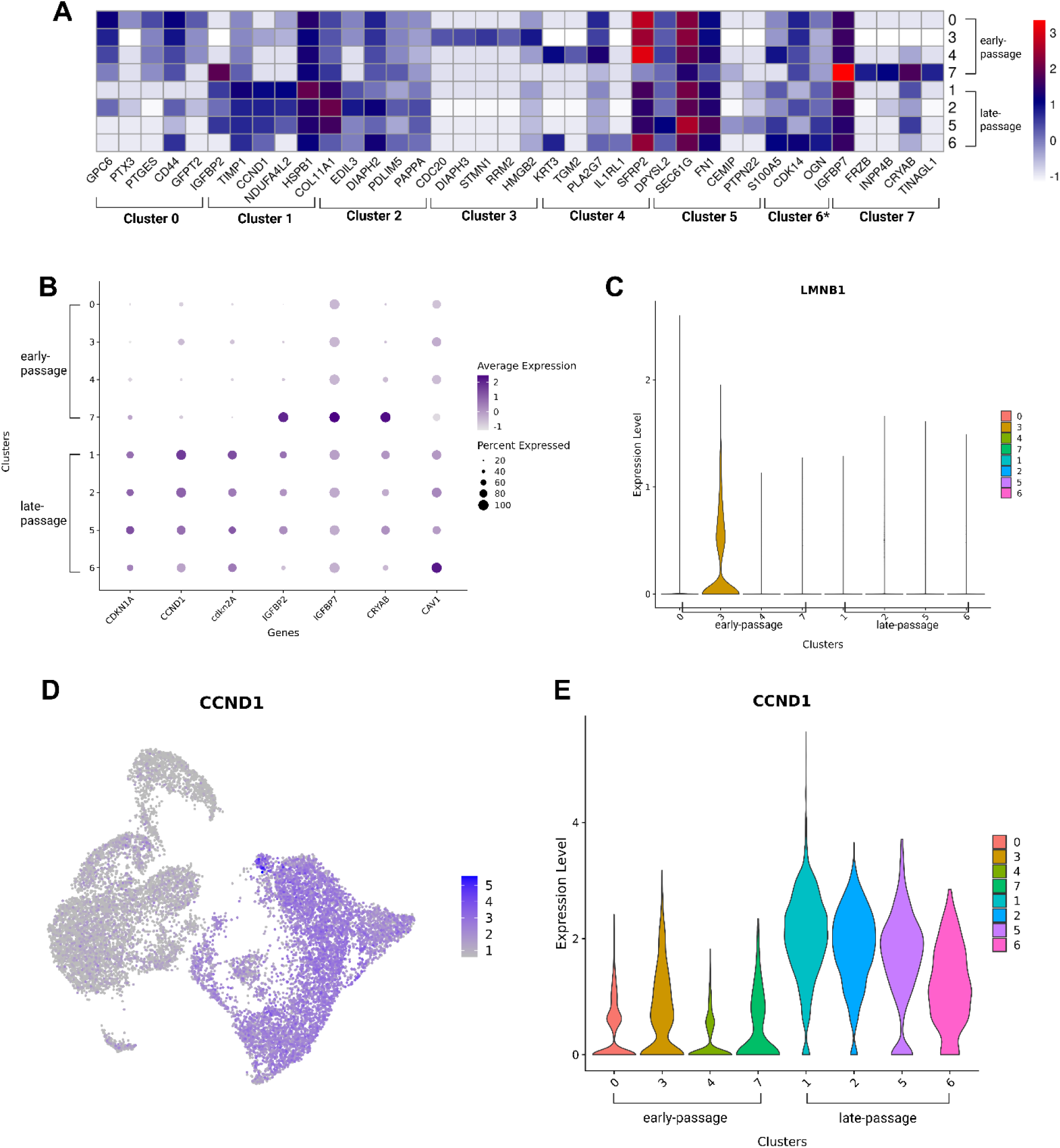
Characterization of cluster-specific markers and assessment of transcriptional patterns of genes associated with cellular senescence. **A.** Heatmap showing the top five highly expressed marker RNAs in each cluster (based on LogFC). Cluster6*: Genes KRT3 and IL1RL1 appear in the top five marker list for both cluster 4 and cluster 6. To avoid redundancy and potential confusion, the expression values of these genes were visualized under cluster 4 and were excluded from the visualization for cluster 6. Median expression levels were plotted for cells in each cluster. Colour represents expression intensity (based on LogFC value). The clusters are arranged based on the origin of the sample (P2 followed by P6), whereas the genes are arranged based on cluster order from cluster 0 to 7. **B.** Dotplot depicting markers commonly associated with cellular senescence, with dot colour representing the average RNA expression levels scaled across all clusters. The size of each dot indicates the percentage of cells expressing a specific RNA within each cluster. The clusters are arranged based on the origin of the sample (P2 followed by P6). **C.** Violin plot depicting the expression of LMNB1. **D.** Feature plot depicting the expression of CCND1. **E.** Violin plot showing the expression level of CCND1 for each cluster. The clusters are arranged based on the origin of the sample (P2 followed by P6).

We further evaluated the expression of several genes commonly associated with the hallmarks of cellular senescence. Notably, senescence-associated genes encoding the proteins p21 (CDKN1A), p16 (CDKN2A), and cyclin D1 (CCND1) were overexpressed in the late-passage cell clusters. The SASP factors IGFBP2 and IGFBP7 displayed increased expression in cluster 7; however, IGFBP2 was downregulated in the other P2-derived clusters, while IGFBP7 demonstrated consistent expression across all remaining clusters. Additionally, the newly identified robust senescence marker CRYAB was also significantly overexpressed in cluster 7.

Furthermore, CAV1, a gene known to play a critical role in cellular senescence, exhibited marked upregulation in cells from cluster 6 (Figure 7B). Loss of LMNB1 was proposed to be associated with cellular senescence[57, 58], and, in line with this, our study revealed a pronounced downregulation of LMNB1 expression in late-passage cell clusters; interestingly, with a similar reduction in early-passage-derived clusters 0, 4, and 7. However, as shown in Figure 7C, LMNB1 expression remained consistently high in cluster 3, a population of actively proliferating cells. Among these identified genes, CCND1 was one of the core genes exhibiting elevated expression levels in all late-passage clusters compared to early-passage clusters (Figures 7D and E).

### Inter-cluster comparison reveals distinct gene expression profiles of late-passage cMSCs

To gain deeper insight into the transcriptomic profiles of late-passage-derived senescent cells and their distribution, we conducted an inter-cluster comparison between early-passage and late-passage cell clusters. Specifically, transcriptional profiles of early-passage clusters (0, 3, 4, 7) were compared to those of late-passage clusters (1, 2, 5, 6) to identify differential gene expression patterns associated with cellular senescence. This approach allowed us to explore the distinct transcriptional signatures potentially contributing to the onset and progression of senescence, providing a clearer understanding of the molecular shifts that occur during cellular aging at the single-cell level. The comparison yielded a total of 3,319 differentially expressed genes; among these, the ten genes exhibiting the highest expression levels (as determined by logFC values), along with the ten most significantly downregulated genes (Supplementary Table 13), are presented in Figure 8A. Commonly recognized markers of cellular senescence including CCND1, IGFBP2, CDKN1A, CDKN2A, and TIMP1 were considerably overexpressed in the late-passage clusters. Moreover, CRYAB, an important senescence-related gene and novel senolytic target[59, 60], also showed elevated expression in the late-passage clusters. Interestingly, CRYAB, along with IGFBP2, presented considerable enrichment in cluster 7, suggesting that the early-passage-derived cells constituting this cluster may also be in a senescent state.

**Figure 8.**
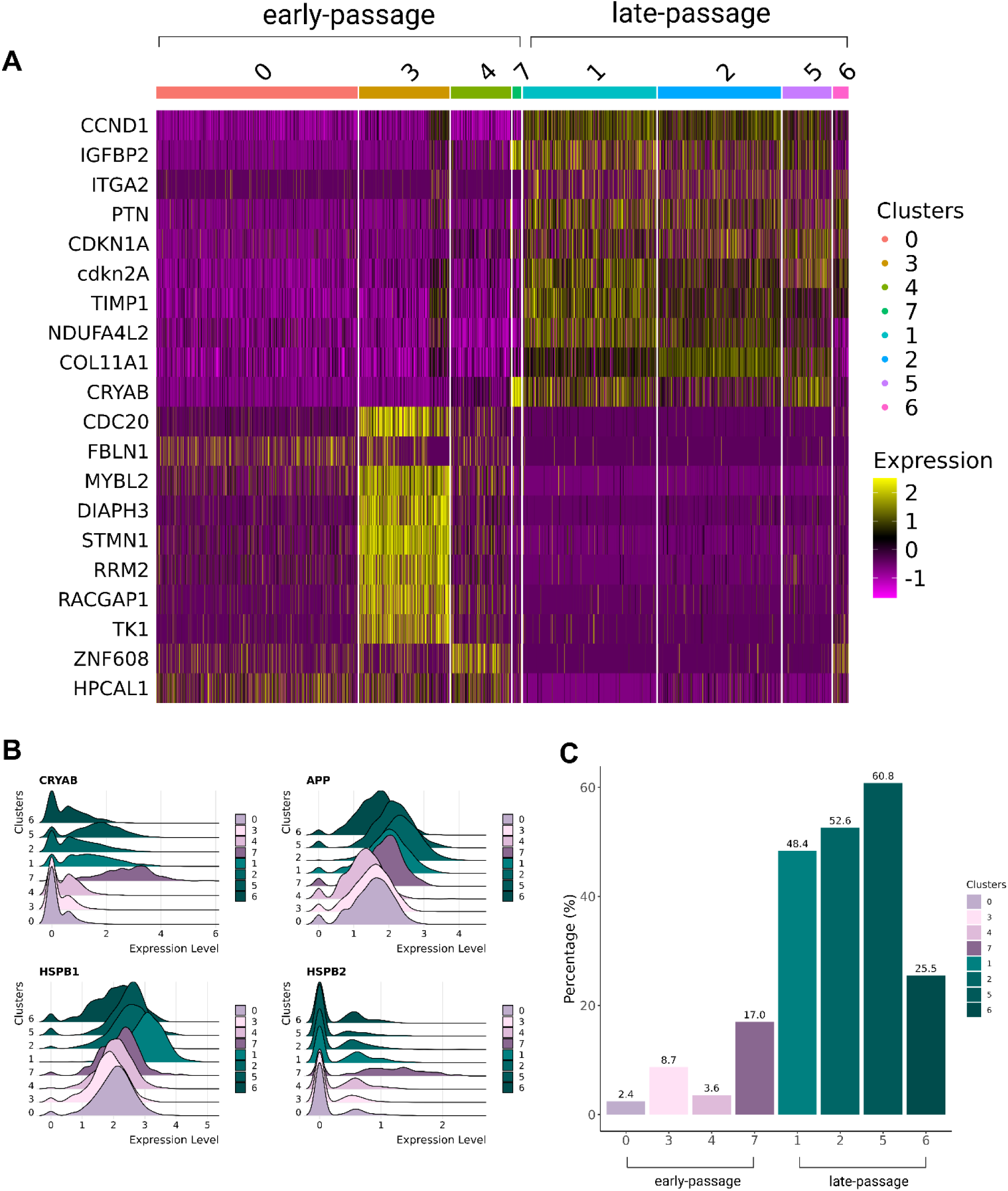
Analysis of top marker genes in late-passage cell populations and characterisation of CRYAB-interacting partners. **A.** Heatmap showing the expression of the top ten marker genes in late-passage cell populations. Each line represents a single cell, the colour indicates the expression level (pink – downregulated, yellow – overexpressed). The clusters are arranged based on the origin of the sample (P2 followed by P6). **B.** Ridge plot of CRYAB expression and its interacting partners APP, HSPB1, and HSPB2. **C.** Bar graph displaying the percentage of putative senescent cells across clusters, based on gene co-occurrence analysis. The analysis includes both established senescence markers and newly identified genes that may serve as potential novel markers of cellular senescence. The clusters are arranged based on the origin of the sample (P2 followed by P6).

We also evaluated the expression levels of the genes most significantly downregulated in P6, identified by their lowest logFC values, and from the top five marker genes of cluster 3, four were among them: CDC20, DIAPH3, STMN1, and RRM2, indicating that cluster 3 also plays an important role in driving the differences between early and late-passage cell populations. Several key genes involved in fundamental cellular processes were also downregulated in the late-passage clusters. These include FBLN1, which promotes osteogenesis[61]; MYBL2, a critical regulator of cell cycle progression; RACGAP1, a gene that regulates cytokinesis, cell growth, and differentiation; TK1, essential for DNA replication; the transcription factor ZNF608; and HPCAL1, which has been shown to promote glioblastoma cell proliferation[62] (Figure 8A). We further investigated the expression profiles of the novel senolytic target, CRYAB and its interacting partners across the cell clusters. As previously highlighted, CRYAB exhibited marked overexpression in the late-passage clusters, but with cluster 7 showing the most pronounced upregulation. Additionally, CRYAB’s interacting partners APP and HSPB1 were also overexpressed in late-passage clusters relative to early-passage ones. Notably, APP demonstrated elevated expression in cluster 2, while HSPB1 showed substantial overexpression in cluster 1.

Interestingly, HSPB2 expression was upregulated in cluster 7, further supporting the senescence-associated phenotype observed in this cluster (Figure 8B).

Building on these results, we aimed to assess our approach by quantifying the proportion of candidate senescent cells within each cluster. Therefore, a gene co-occurrence analysis was performed under the hypothesis that a cell must express core senescence markers – specifically CCND1, IGFBP2, CDKN1A, and TGFB2 – alongside at least one of the top ten genes (CDKN2A, IGFBP7, CRYAB, ITGA2, PTN, NDUFA4L2, COL11A1) upregulated in P6 as identified through inter-cluster comparison of early-and late-passage cell clusters. The gene co-occurrence analysis revealed a pronounced increase in the proportion of putative senescent cells within late-passage clusters compared to early-passage ones (Figure 8C). Specifically, clusters 0, 3, 4, and 7 in early-passage cells contained 2.4%, 8.7%, 3.6%, and 17% senescent cells, respectively. In contrast, the late-passage clusters 1, 2, 5, and 6 showed markedly higher proportions of these cells, with 48.4%, 52.6%, 60.8%, and 25.5%, respectively (Supplementary Table 14).

### Comparative analysis of senescent and actively dividing subpopulations reveals distinct proliferative and secretory signatures

To further examine the heterogeneity among senescent subpopulations, we focused on clusters 3 and 7, which differ in both cellular composition and senescence-associated gene expression.

Cluster 3 contains a mixture of early-and late-passage cells, whereas cluster 7 is composed primarily of early-passage cells that nevertheless display pronounced SASP activity and senescence signatures. To elucidate the molecular distinctions between these cell groups and their relationship to senescence progression, we performed an inter-cluster comparison. This analysis revealed that cluster 3 exhibits strong expression of proliferation-associated genes (e.g., MYBL2, RRM2, HMGB2, CDC20, STMN1), consistent with an actively dividing population. In contrast, cluster 7 showed marked upregulation of CRYAB, IGFBP7, FRZB, DCN, and SASP-and extracellular matrix-related genes, along with decreased expression of proliferative markers, features characteristic of a senescent, secretory phenotype (Supplementary Figure 8).

Furthermore, to gain a deeper understanding of the mixed composition of cluster 3, which contained both early-and late-passage cells, we conducted a subclustering analysis. UMAP visualization revealed that cluster 3 could be further subdivided based on passage number and the expression of proliferation-related genes. Cells from passage 2 and passage 6 formed largely distinct subgroups with minimal overlap, indicating that passage number is a major contributor to the heterogeneity observed within cluster 3 (Supplementary Figure 9A-D). Within these subclusters, the differential expression of proliferation-associated genes PLK1, MKI67, and MCM3 was evident. Certain subpopulations exhibited high expression of these markers, whereas others showed lower expression levels, suggesting reduced proliferative activity. This refined subclustering thus underscores the intrinsic heterogeneity within cluster 3 (Supplementary Figures 9A-E).

### Evaluation of inter-donor and inter-species consistency of the senescence-related characteristics and identified potential marker genes

To further assess inter-donor consistency and characterize cellular phenotypes, we conducted additional analyses on a second canine isolate, including morphological examination, BrdU incorporation to evaluate proliferative capacity, and single-cell RNA sequencing combined with cell cycle analysis. Similarly to the primary samples, this isolate exhibited a marked decline in proliferation, decreasing from 84% at passage 2 to 56% at passage 6 (Supplementary Figures 10A and 10B), accompanied by an enlarged, flattened morphology characteristic of senescent cells (Supplementary Figure 11). Overall, both isolates demonstrated a pronounced reduction in replicative potential by passage 6, reflecting consistent senescence-associated phenotypes across donors.

The transcriptomic profile was further examined using single-cell sequencing. UMAP clustering revealed ten distinct cellular clusters (Supplementary Figure 12A). When coloured by sample origin (Supplementary Figure 12B), cells from passage 2 and passage 6 showed partial separation, indicating some transcriptional divergence between passages, though mixed clusters were also observed, suggesting overlap in cell states. Cell cycle phase analysis was also conducted and further demonstrated differences in the distribution of cells across G1, S, and G2/M phases between the two passages, with a pronounced enrichment of passage 6 cells in the G1 phase (76.38%) compared to passage 2 cells (43.50%) (Supplementary Figures 12 C and D). To minimize technical variation between samples, the cells from both single-cell RNA sequencing datasets were integrated for downstream analysis. Following integration, the UMAP visualization (Supplementary Figure 13A) revealed that cells from passage 2 and passage 6 each formed distinct and well-defined clusters, with cells from the two sequencing runs of the same passage grouping together. This indicates strong consistency between replicates. A limited overlap was observed between passages, which originated from the second canine isolate. Clustering analysis (Supplementary Figure 13B) identified nine major cell populations, and the distribution of cells across clusters (Supplementary Figure 13C) confirmed this observation, highlighting transcriptomic differences between passages while showing reproducibility across sequencing runs and between different donors. Next, the marker genes identified in the present study were evaluated in the integrated dataset. As illustrated, the expression patterns of the downregulated genes (Supplementary Figure 13D) CDC20, FBLN1, MYBL2, DIAPH3, STMN1, RRM2, RACGAP1, TK1, ZNF608, and HPCAL1 and upregulated genes (Supplementary Figure 13E) CCND1, IGFBP2, CDKN1A, TGFB2, CDKN2A, IGFBP7, CRYAB, ITGA2, PTN, NDUFA4L2, COL11A1, and TIMP1 were consistent with those observed in the first single-cell sequencing run. Specifically, the downregulated genes showed diminished levels, while the upregulated genes showed elevated expression levels in the cell populations.

These consistent expression trends across independent sequencing runs and independent canine isolates further support the inter-donor reproducibility and robustness of the findings.

To evaluate interspecies consistency of the identified candidate markers, we performed a literature search to determine publicly available datasets suitable for comparison. We specifically sought studies aligning with our experimental context of replicative senescence. Accordingly, publicly available mouse and human replicative exhaustion datasets were analysed to assess the expression patterns of the proposed marker genes. Differential gene expression analysis of the mouse dataset, comparing late-passage (P7) to early-passage (P1) cells, demonstrated that CCND1, IGFBP2, CDKN1A, CDKN2A, IGFBP7, PTN, NDUFA4L2, and TIMP1 were significantly upregulated in P7 cells, consistent with the described senescence-associated transcriptional profile. In contrast, ITGA2 showed downregulation, while TGFB2 and CRYAB were not detected among the differentially expressed genes. Examination of the downregulated markers revealed that CDC20, MYBL2, DIAPH3, STMN1, RRM2, RACGAP1, and TK1 were also significantly reduced in expression, whereas FBLN1, ZNF608, and HPCAL1 did not show significant differential expression between passages (Supplementary Table 15).

Furthermore, differential gene expression analysis of the human dataset, comparing proliferative cells to cells undergoing replicative senescence, revealed that CCND1, TGFB2, IGFBP7, and ITGA2 were upregulated, whereas PTN, NDUFA4L2, and COL11A1 were downregulated.

Among the proposed downregulated markers, FBLN1, MYBL2, STMN1, RRM2, and TK1 also showed decreased expression, while HPCAL1 was the only gene displaying increased expression in senescent cells (Supplementary Table 16).

Comparative analysis across these species revealed a total of nine genes that were significantly differentially expressed in dog, human, and mouse datasets. Among these, six genes displayed consistent expression patterns (either upregulated or downregulated) across all three species, indicating strong interspecies conservation of senescence-associated transcriptional responses. Additionally, nine genes were shared between the dog and mouse datasets, suggesting a high degree of overlap in their senescence-related gene expression profiles. Detailed gene lists and overlap are provided in Supplementary Figure 14.

In addition, single-cell RNA sequencing data from a recent study by Taherian Fard et al. (2024)[48] investigating human MSC replicative senescence were analysed independently. We selected all replicates corresponding to two time points: T0 (day 23) representing proliferative cells and T2 (day 63) representing cells undergoing replicative senescence. This comparison identified 16 genes exhibiting expression patterns consistent with our dataset, with CCND1, CDKN1A, CDKN2A, CRYAB, ITGA2, NDUFA4L2, COL11A1, and TIMP1 showing upregulation (Supplementary Figure 15A), while CDC20, FBLN1, MYBL2, DIAPH3, STMN1, RRM2, RACGAP1, and TK1 were markedly downregulated in T2 samples (Supplementary Figure 15B).

## Discussion

In the rapidly advancing field of cell therapy, MSCs have captured considerable interest due to their unique biological properties, stemness, and immunomodulatory abilities[63, 64].

Nevertheless, their efficacy can diminish due to replicative senescence. While a variety of biomarkers and techniques have been explored to identify and characterize senescent MSCs, the lack of a universally accepted, specific marker complicates reliable detection and characterisation of these cells. Our study contributes to solving this issue by providing a detailed analysis of the transcriptional changes associated with replicative senescence of MSCs in dog, a model organism known for its key physiological similarities to humans and recognized as a valuable platform for aging research[25],[65–67].

cMSCs exhibit both conserved and species-specific characteristics of senescence, reflected by fundamental differences in telomere biology and cellular stress responses. In humans, replicative senescence is characterized by enlarged, flattened cell morphology, strong SA-β-gal accumulation, and upregulation of p16 and p21, accompanied by a broad transcriptional shift toward a SASP-related phenotype[28]. On the other hand, murine MSCs may exhibit similar morphological and β-gal changes but progress more slowly toward senescence, largely due to their long telomeres and sustained telomerase activity[68]. Moreover, Fehrer et al. (2006) observed that murine MSCs lack the classical indicators of *in vitro* senescence, showing no growth arrest or SA-β-gal activity, although cells with abnormal morphology emerge at higher passages[12]. However, studies have shown that the transcript for p16, encoded by the CDKN2A gene, exhibits a similar age-associated increase in expression in humans and rodents[69], [70]. cMSCs exhibit classical features of senescence similar to human MSCs, including altered morphology, reduced proliferation, and increased β-galactosidase activity[71] and also tend to enter senescence more rapidly[72]. However, studies examining gene expression changes during cMSC senescence remain limited. Therefore, cMSCs closely reflect human senescence and support longitudinal studies in a large-animal system, positioning the dog as a highly relevant model for investigating MSC senescence. In this study, we employed a passage-based *in vitro* approach of replicative senescence by culturing cMSCs until they reached replicative exhaustion. To this end, we compared early passage P2 and late passage P6 cells exhibiting markedly distinct proliferative capacities, as clearly reflected by their respective PDLs.

Replicative senescence in cMSCs was first indicated by distinct morphological alterations, with late-passage P6 cells exhibiting the characteristic enlarged and flattened morphology, elevated SA-β-gal activity, and significantly reduced BrdU incorporation, confirming the senescent phenotype, consistent with previously reported senescent phenotypes[73, 74]. Employing bulk RNA sequencing to assess gene expression changes associated with replicative senescence, we identified a set of significantly upregulated and downregulated genes in P6 cells compared to early-passage P2 cells. Among the significantly upregulated genes, we detected well-established senescence markers, including CCND1, CDKN1A, and CDKN2A. Moreover, we identified genes with the highest levels of overexpression in P6 MSCs, some of which correspond to known senescence-associated signatures, while others point to potentially novel associations. For instance, VCAM1, which was previously described as being overexpressed in senescent endothelial cells[75, 76], was significantly upregulated in our study. Among other notable genes, ANKRD1 has been reported as aging-related[77] and senescence marker gene[78], while MECOM has been associated with pathways relevant to senescence, cell cycle, and p53 signalling, as highlighted by a former study by KEGG analysis[79]. Moreover, it is important to highlight that other genes such as NKX2-5, LHX1, ADGRB1, and CPA4 displayed significant upregulation but lack prior associations with cellular senescence. Additionally, TIE1, shown to exhibit age-independent upregulation in vascular contexts[80], and HOXA13, described as a key participant in Erk1/2 activation[81, 82], may offer unique insights into cellular aging mechanisms, while NEFH, a biomarker of neuronal differentiation[83], which demonstrated the highest overexpression, and C1QL1, a senescence-associated gene with potential roles in neuronal differentiation[84, 85], suggest potential new directions for exploration.

Furthermore, numerous genes linked to cell proliferation and cell cycle regulation appear markedly downregulated in senescent P6 cells compared to early-passage P2 cells. Among the most significantly downregulated were BNC1, a transcription factor predominantly expressed in proliferative keratinocytes and germ cells[86]; SELENOP, which plays a role in Wnt pathway activation and cellular proliferation[87]; LSP1, involved in the negative regulation of proliferation[88]; and PREX1, which supports homeostatic proliferation[89]. Several other genes including critical regulators of cell proliferation and cell cycle progression were also observed to be downregulated in the senescent P6 samples, namely, TOP2A, MYBL2, MKI67, DPT, WT1, and PIMREG[90–95].

Another defining feature of senescent cells is stable cell cycle arrest, characterized by the loss of proliferative capacity. The downregulation of these genes potentially facilitates the advancement of cellular senescence. The loss of LMNB1, a critical constituent of the nuclear lamina, has been shown to represent another important characteristic and marker of senescent cells[16]. Consistent with this, our results showed a clear and significant downregulation of this gene in the senescent P6 samples.

Moreover, GO and KEGG enrichment analyses provided deeper insight into the biological relevance of the identified DEGs in the context of senescence. Downregulated genes were primarily associated with cell division and cell cycle-related processes – such as mitotic spindle assembly checkpoint signalling, G2/M phase transition, and DNA replication – highlighting a functional shift away from proliferation. Additional GO terms, such as regulation of the mitotic cell cycle, mitotic checkpoint signalling, and cell cycle regulation, further emphasize this pattern. These findings are consistent with prior research demonstrating that senescent cells commonly undergo stable cell cycle arrest and reduced proliferative activity[96–98]. This evidence further supports the hypothesis that late-passage P6 samples are not actively dividing and are likely in a state of cell cycle arrest.

The analysis also highlighted significantly enriched GO terms for the upregulated DEGs corresponding to biological processes closely related to and previously associated with aging and cellular senescence, such as the ERK1 and ERK2 cascade[99], the integrin-mediated signalling pathway[100], and response to hypoxia[101]. Moreover, the KEGG pathway analysis identified key pathways that are actively studied in the context of cellular senescence, such as the HIF-1 signalling pathway, which is essential for response to hypoxia[102], and the PI3K-Akt signalling pathway[103]. These pathways are well-established in the literature as key players in cellular aging and senescence, often contributing to both cell survival and the pro-inflammatory features of senescence. These findings suggest that senescence is not merely a state of arrest, but an actively regulated process with significant implications for tissue homeostasis and aging.

However, during replicative senescence, cells with critically shortened telomeres trigger a DNA damage response resulting in cell cycle arrest[104]. The cyclin-dependent kinase inhibitors p21 and p16 play crucial roles in this process by orchestrating proliferative arrest. These proteins have been widely recognized as hallmark markers of senescent cells[105]. In our study, flow cytometry using propidium iodide labelling, followed by single-cell analysis with CellCycleScoring, revealed a pronounced accumulation of P6 cells in the G1 phase. This approach provided a more sensitive and precise identification of cell cycle phases, utilizing gene expression markers of proliferation. Furthermore, the expressions of the proliferation-and cell division-specific markers MCM3, MCM6, MKI67, and PLK1 were assessed for the clusters. All these markers were found to be downregulated in the P6 clusters. Interestingly, among P2 clusters, only cluster 3 showed marked expression of MCM3, MKI67, and PLK1, further supporting that P6-derived cells are no longer capable of sustained division, while early-passage cells, such as those in cluster 3, remain proliferatively active.

The development of single-cell sequencing technologies enabled us to uncover that the gene expression changes identified in our study demonstrate considerable heterogeneity at the level of individual cells. Therefore, the expressions of genes associated with important identified GO terms – representing pathways involved in senescence, including the ERK1/ERK2 signalling cascade, the integrin-mediated signalling pathway, and the cellular response to oxygen levels – were also evaluated at the single-cell level. In accordance with bulk RNA sequencing results, late-passage cell populations from clusters 1, 2, 5, and 6 expressed elevated levels of APP, EDN1, and CCL5. However, cluster 6 demonstrated reduced expression of EDN1 and CCL5 compared to the other P6-derived clusters, suggesting potential differences in the activation of the ERK1/ERK2 pathway within specific subpopulations. The integrin-mediated signalling pathway, another significantly enriched GO term in P6 cells, showed upregulation of key genes, including TIMP1, ITGA1, and ITGA2, across late-passage clusters. Interestingly, ITGA1 was more highly expressed in cluster 7, while ITGA2 was prominently expressed in clusters 2 and 6. Furthermore, genes involved in the cellular response to oxygen levels, a critical factor in senescence induction[106], were generally upregulated in P6-derived clusters. However, similarly to the pattern observed in the ERK1/ERK2 pathway, EDN1 – also included in the oxygen levels GO term – was downregulated in cluster 6.

Kawamura et al. recently reported upregulation of TGFB2 in old MSCs compared to younger MSCs[107]. Similarly, our study identified a marked upregulation of TGFB2 across all clusters derived from late-passage MSCs, further supporting these observations. Furthermore, Caveolin 1 (CAV1), a key regulator of cellular senescence[108], which has also been shown to be upregulated in response to oxidative stress[60], exhibited significant upregulation in cluster 6.

CRYAB also exhibited elevated expression in all P6 clusters compared to P2; intriguingly, it also showed particularly high expression in cluster 7 of the P2 population. This finding further supports the involvement of oxygen-related stress in senescence, as CRYAB is known to be upregulated in response to oxidative stress conditions[109]. Collectively, these findings underscore the complexity and heterogeneity of cellular senescence, highlighting the distinct activation of senescence-related pathways across different late-passage cell populations.

UMAP analysis of the dataset from P2-and P6-derived cells revealed eight transcriptionally distinct clusters. Visualization by cell origin demonstrated near-complete segregation between P2 and P6 populations; however, cluster 3 stands out as less well-defined, consisting predominantly of P2-derived cells but also including a notable subset from P6, suggesting partial overlap in gene expression profiles. Differences in transcriptional profiles were also observed among the identified clusters. To characterize the clusters based on their expression profiles, we employed the ‘FindAllMarkers’ function to identify genes that were most prominently expressed in each cluster. This approach revealed considerable variation in the transcriptional patterns across the clusters, offering important insights into their unique gene expression signatures. Notably, cluster 3 exhibited a distinct transcriptional profile, characterized by the expression of genes that play crucial roles in cellular processes, including those involved in cell division and proliferation.

This suggests that cluster 3 comprises primarily actively dividing cells, while also containing a subset of active cells from P6, further highlighting its role as a proliferative population. In contrast, clusters associated with late-passage P6 cells exhibited a high representation of key senescence and SASP-related genes, including TIMP1, CRYAB, CCND1, IGFBP2, IGFBP7, and PAPPA. Interestingly, cluster 7, derived from P2 cells, displayed the most pronounced expression of IGFBP7 and CRYAB.

The induction of cell cycle arrest and the onset of senescence were corroborated by evaluating the expression of the cyclin-dependent kinase inhibitors p21 (CDKN1A) and p16 (CDKN2A), which play critical roles in regulating proliferative arrest at the single-cell level. CCND1 is well known to promote progression through the G1/S checkpoint, and its overexpression can stimulate cell cycle progression or bypass arrest[110]. However, in certain contexts, CCND1 overexpression can paradoxically induce cell cycle arrest or senescence-like phenotypes.

Sustained CCND1 accumulation may result in aberrant CDK activity, activate checkpoint pathways such as the p21 signalling cascade, or disrupt regulatory feedback, ultimately reinforcing growth arrest rather than promoting proliferation[111]. Additionally, acute overexpression of cyclin D1 (CCND1) has been demonstrated to drive cell cycle arrest in the G1 phase[112]. However, previous studies showed downregulation of CCND1 in senescent human MSCs[113] and that its expression can counteract senescence, whereas loss of CCND1 accelerates senescence in certain cancer cell models[114]. This apparent discrepancy may reflect context-or tissue-dependent regulation, which may account for the observed upregulation of CCND1 in senescent cMSCs, despite their persistent G1 arrest.

Our results showed a considerable overexpression of the CDKN1A and CDKN2A genes across all P6-derived clusters, as compared to P2 cell clusters. Surprisingly, SASP factors, such as IGFBP2, IGFBP7, and CRYAB, which were described as an oxidative stress-related genes[109] and potential senolytic target[59], exhibited substantial overexpression specifically in cluster 7. The presence of these markers in a cluster originating from early-passage cells suggests that a subset of P2 cells may already be undergoing stress responses or entering a pre-senescent state. This observation points to a potential heterogeneity in senescence susceptibility within cluster 7, originating from early-passage populations. Interestingly, we observed elevated OGN expression across all late-passage cMSC clusters, despite previous reports of OGN downregulation in senescent human MSCs[115] and its link to reduced osteogenic capacity in aged MSCs[116].

This upregulation may reflect a compensatory mechanism in cMSCs to preserve osteogenic potential during senescence, also highlighting possible species-or context-dependent differences in OGN regulation. Additionally, previous studies have described that the loss of LMNB1, a key component of the nuclear lamina, is a prominent marker of senescent cells[16]. In line with our bulk RNA sequencing results, P6 samples exhibited a clear downregulation of LMNB1.

Interestingly, within the P2 cell populations, only cluster 3 displayed marked expression of LMNB1, a finding that further supports the proliferatively active status of the cells in this cluster. Interestingly, while LMNB1 expression was downregulated across P6-derived cells, as expected due to its known downregulation during senescence, its lack of pronounced expression in cluster 7 compared to cluster 3 further suggests that cluster 7 may be diverging from the proliferative state and adopting a distinct phenotype. This intermediate or transitional state between proliferation and senescence is particularly intriguing and may have significant implications for understanding the onset of cellular senescence.

In late-passage cells, inter-cluster comparison revealed upregulation of established senescence markers (CCND1, CDKN1A, CDKN2A) alongside genes previously linked to senescence (IGFBP2, TGFB2, IGFBP7, ITGA2, PTN, COL11A1, TIMP1, CRYAB), while NDUFA4L2 appeared as a potentially novel association. Among the downregulated genes, FBLN1, DIAPH3, and ZNF608 represent previously unreported candidates that may have unexplored roles in the process of cellular senescence. We also showed that CRYAB’s interacting partners were upregulated in P6 clusters, except for HSPB2, which exhibited particularly notable overexpression in cluster 7, providing additional evidence that cluster 7 might also present a senescence-related phenotype. Comparative analysis of senescent and proliferative subpopulations revealed distinct transcriptional patterns underlying their divergent phenotypes. Cluster 3, composed of both early-and late-passage cells, showed strong expression of proliferation-associated genes (e.g., MYBL2, RRM2, HMGB2, CDC20, STMN1), indicating that a subset of late-passage cells also retained proliferative activity. Moreover, subclustering of cluster 3 revealed that passage number was a key driver of intracluster heterogeneity, with distinct subgroups emerging based on differential expression of proliferative markers. In contrast, cluster 7 was characterized by marked upregulation of CRYAB, IGFBP7, and other SASP-and ECM-related genes, reflecting a senescent, secretory phenotype. Notably, the presence of senescence-associated markers in early-passage cluster 7 suggests that pre-senescent features can emerge prior to replicative exhaustion, highlighting the intrinsic heterogeneity and gradual onset of senescence within MSC populations.

To evaluate the reproducibility and biological relevance of these findings, we performed cross-species comparisons using publicly available human and murine MSC senescence datasets. This analysis revealed strong conservation of transcriptional trends across independent canine isolates and consistency among species, confirming the robustness of the proposed marker panel. Gene co-occurrence analysis further demonstrated a substantially higher proportion of senescent cells in late-passage clusters, validating the transcriptional indicators identified.

Beyond providing a molecular map of MSC senescence, our findings have important translational implications. The identified gene panel could guide the development of platforms for monitoring MSC quality in therapeutic manufacturing. In particular, these markers could be integrated into prospective quality control assays to detect early transcriptional indicators of senescence before the functional decline occurs. However, translating transcriptomic signatures into practical diagnostic assays will require further validation. Overall, this study provides a comprehensive characterization of replicative senescence in cMSCs, revealing distinct molecular states that capture the transition from proliferation to senescence. By defining both conserved and novel markers and proposing their combined use as a panel of up-and downregulated genes for MSC manufacturing and quality control, our findings bridge the gap between mechanistic discovery and clinical translation, offering a foundation for improved standardization and efficacy in MSC-based therapies.

## List of abbreviations

DNA: Deoxyribonucleic acid
RNA: Ribonucleic acid
S phase: Synthesis phase
MSC: Mesenchymal Stem Cell
cMSC: Canine mesenchymal stem cell
hMSC: Human Mesenchymal Stem Cell
P2: Passage 2
P6: Passage 6
SA-β-galactosidase: Senescence-associated-β-galactosidase
PD: Population doubling
SASP: Senescence-associated secretory phenotype
IQR: Interquartile range
PCA: Principal Component Analysis
DEG: Differentially expressed gene
NGS: Next-generation sequencing
TPM: Transcripts Per Million
UMAP: Uniform Manifold Approximation and Projection

## Supplementary information

All data generated or analysed during this study are included in this published article and its supplementary information files: Supplementary information and Supplementary Data.

## Declarations

### Ethics Approval and Consent to Participate

The owner of the dogs who participated in this study provided written informed consent.

## Consent for publication

Not applicable.

## Availability of data and materials

The RNA-sequencing and single-cell RNA sequencing data generated in this study have been deposited in the NCBI Sequence Read Archive (SRA) under the BioProject accession numbers PRJNA1235683 and PRJNA1235986, respectively. The data for the second single-cell RNA sequencing run are available from the following link: https://doi.org/10.5281/zenodo.17346986.

The code utilized to analyse the data and to generate the figures presented in this study are available in the GitHub repository based on the following link: <u>GitHub-erdaqorri/Mesenchymal-Stem-Cell-CLF</u>

## Declaration of generative AI and AI-assisted technologies in the writing process

During the preparation of this work the author(s) used ChatGPT-4o in order to enhance the language quality of the manuscript. After using this tool, the author(s) reviewed and edited the content as needed and take(s) full responsibility for the content of the publication.

## Authors’ contributions

EP contributed to the conceptualization, methodology, investigation, formal analysis, and writing – original draft manuscript, visualization. EQ contributed to methodology, bioinformatical analysis, visualization, formal analysis, and writing – original draft manuscript. MZE contributed to methodology, writing, and editing. VS contributed to methodology, writing, and editing. FA contributed to methodology, writing, and editing, ÉSK contributed to methodology. CB contributed to methodology and formal analysis. MM contributed to methodology and formal analysis. FS contributed to formal analysis and resources. EKT contributed to conceptualization, methodology, and formal analysis. LH contributed to conceptualization, formal analysis, project administration, resources, and funding acquisition. All authors read and approved the final manuscript.

## Acknowledgements

We would like to sincerely thank Gabriella Tick for her contribution in proofreading the manuscript.

We thank Edit Kotogány and the Laboratory of Functional Genomics at the HUN-REN Biological Research Centre, Szeged, for the support in performing the flow cytometry analysis.

## References

1. El Omar R, Beroud J, Stoltz J-F, Menu P, Velot E, Decot V. Umbilical Cord Mesenchymal Stem Cells: The New Gold Standard for Mesenchymal Stem Cell-Based Therapies? Tissue Eng Part B Rev. 2014;20:523–44. 10.1089/ten.teb.2013.0664.

2. Abbaszadeh H, Ghorbani F, Derakhshani M, Movassaghpour AA, Yousefi M, Talebi M, et al. Regenerative potential of Wharton’s jelly-derived mesenchymal stem cells: A new horizon of stem cell therapy. J Cell Physiol. 2020;235:9230–40. 10.1002/jcp.29810.

3. Lindner U, Kramer J, Rohwedel J, Schlenke P. Mesenchymal Stem or Stromal Cells: Toward a Better Understanding of Their Biology? Transfusion Medicine and Hemotherapy. 2010;37:75–83. 10.1159/000290897.

4. Wong P-F, Dharmani M, Ramasamy TS. Senotherapeutics for mesenchymal stem cell senescence and rejuvenation. Drug Discov Today. 2023;28:103424. 10.1016/j.drudis.2022.103424.

5. Oh J, Lee YD, Wagers AJ. Stem cell aging: mechanisms, regulators and therapeutic opportunities. Nat Med. 2014;20:870–80. 10.1038/nm.3651.

6. Goodell MA, Rando TA. Stem cells and healthy aging. Science (1979). 2015;350:1199–204. 10.1126/science.aab3388.

7. Carlos Sepúlveda J, Tomé M, Eugenia Fernández M, Delgado M, Campisi J, Bernad A, et al. Cell Senescence Abrogates the Therapeutic Potential of Human Mesenchymal Stem Cells in the Lethal Endotoxemia Model. Stem Cells. 2014;32:1865–77. 10.1002/stem.1654.

8. Liu J, Ding Y, Liu Z, Liang X. Senescence in Mesenchymal Stem Cells: Functional Alterations, Molecular Mechanisms, and Rejuvenation Strategies. Front Cell Dev Biol. 2020;8. 10.3389/fcell.2020.00258.

9. Sethe S, Scutt A, Stolzing A. Aging of mesenchymal stem cells. Ageing Res Rev. 2006;5:91–116. 10.1016/j.arr.2005.10.001.

10. Prišlin M, Butorac A, Bertoša R, Kunić V, Ljolje I, Kostešić P, et al. In vitro aging alters the gene expression and secretome composition of canine adipose-derived mesenchymal stem cells. Front Vet Sci. 2024;11. 10.3389/fvets.2024.1387174.

11. Amit M, Carpenter MK, Inokuma MS, Chiu C-P, Harris CP, Waknitz MA, et al. Clonally Derived Human Embryonic Stem Cell Lines Maintain Pluripotency and Proliferative Potential for Prolonged Periods of Culture. Dev Biol. 2000;227:271–8. 10.1006/dbio.2000.9912.

12. Fehrer C, Laschober G, Lepperdinger G. Aging of murine mesenchymal stem cells. In: Annals of the New York Academy of Sciences. 2006. 10.1196/annals.1354.030.

13. Bruder SP, Jaiswal N, Haynesworth SE. Growth kinetics, self-renewal, and the osteogenic potential of purified human mesenchymal stem cells during extensive subcultivation and following cryopreservation. J Cell Biochem. 1997;64:278–94. 10.1002/(SICI)1097-4644(199702)64:2<278::AID-JCB11>3.0.CO;2-F.

14. Stenderup K. Aging is associated with decreased maximal life span and accelerated senescence of bone marrow stromal cells,. Bone. 2003;33:919–26. 10.1016/j.bone.2003.07.005.

15. Trabucco SE, Zhang H. Finding Shangri-La: Limiting the Impact of Senescence on Aging. Cell Stem Cell. 2016;18:305–6. 10.1016/j.stem.2016.02.002.

16. Freund A, Laberge R-M, Demaria M, Campisi J. Lamin B1 loss is a senescence-associated biomarker. Mol Biol Cell. 2012;23:2066–75. 10.1091/mbc.e11-10-0884.

17. Gardner SE, Humphry M, Bennett MR, Clarke MCH. Senescent Vascular Smooth Muscle Cells Drive Inflammation Through an Interleukin-1α–Dependent Senescence-Associated Secretory Phenotype. Arterioscler Thromb Vasc Biol. 2015;35:1963–74. 10.1161/ATVBAHA.115.305896.

18. Acosta JC, Banito A, Wuestefeld T, Georgilis A, Janich P, Morton JP, et al. A complex secretory program orchestrated by the inflammasome controls paracrine senescence. Nat Cell Biol. 2013;15:978–90. 10.1038/ncb2784.

19. Özcan S, Alessio N, Acar MB, Mert E, Omerli F, Peluso G, et al. Unbiased analysis of senescence associated secretory phenotype (SASP) to identify common components following different genotoxic stresses. Aging. 2016;8:1316–29. 10.18632/aging.100971.

20. Demaria M, Ohtani N, Youssef SA, Rodier F, Toussaint W, Mitchell JR, et al. An Essential Role for Senescent Cells in Optimal Wound Healing through Secretion of PDGF-AA. Dev Cell. 2014;31:722–33. 10.1016/j.devcel.2014.11.012.

21. Coppé J-P, Desprez P-Y, Krtolica A, Campisi J. The Senescence-Associated Secretory Phenotype: The Dark Side of Tumor Suppression. Annual Review of Pathology: Mechanisms of Disease. 2010;5:99–118. 10.1146/annurev-pathol-121808-102144.

22. Biran A, Zada L, Abou Karam P, Vadai E, Roitman L, Ovadya Y, et al. Quantitative identification of senescent cells in aging and disease. Aging Cell. 2017;16:661–71. 10.1111/acel.12592.

23. Schafer MJ, Zhang X, Kumar A, Atkinson EJ, Zhu Y, Jachim S, et al. The senescence-associated secretome as an indicator of age and medical risk. JCI Insight. 2020;5. 10.1172/jci.insight.133668.

24. McKenzie BA. Comparative veterinary geroscience: mechanism of molecular, cellular, and tissue aging in humans, laboratory animal models, and companion dogs and cats. Am J Vet Res. 2022;83. 10.2460/ajvr.22.02.0027.

25. Sándor S, Kubinyi E. Genetic Pathways of Aging and Their Relevance in the Dog as a Natural Model of Human Aging. Front Genet. 2019;10. 10.3389/fgene.2019.00948.

26. Bruder SP, Jaiswal N, Haynesworth SE. Growth kinetics, self-renewal, and the osteogenic potential of purified human mesenchymal stem cells during extensive subcultivation and following cryopreservation. J Cell Biochem. 1997;64:278–94. 10.1002/(sici)1097-4644(199702)64:2<278::aid-jcb11>3.0.co;2-f.

27. Lu LL, Liu YJ, Yang SG, Zhao QJ, Wang X, Gong W, et al. Isolation and characterization of human umbilical cord mesenchymal stem cells with hematopoiesis-supportive function and other potentials. Haematologica. 2006;91.

28. Turinetto V, Vitale E, Giachino C. Senescence in Human Mesenchymal Stem Cells: Functional Changes and Implications in Stem Cell-Based Therapy. Int J Mol Sci. 2016;17:1164. 10.3390/ijms17071164.

29. Khademi-Shirvan M, Ghorbaninejad M, Hosseini S, Baghaban Eslaminejad M. The Importance of Stem Cell Senescence in Regenerative Medicine. 2020. p. 87–102. 10.1007/5584_2020_489.

30. Bertolo A, Baur M, Guerrero J, Pötzel T, Stoyanov J. Autofluorescence is a Reliable in vitro Marker of Cellular Senescence in Human Mesenchymal Stromal Cells. Sci Rep. 2019;9:2074. 10.1038/s41598-019-38546-2.

31. Debacq-Chainiaux F, Erusalimsky JD, Campisi J, Toussaint O. Protocols to detect senescence-associated beta-galactosidase (SA-βgal) activity, a biomarker of senescent cells in culture and in vivo. Nat Protoc. 2009;4:1798–806. 10.1038/nprot.2009.191.

32. Cohn RL, Gasek NS, Kuchel GA, Xu M. The heterogeneity of cellular senescence: insights at the single-cell level. Trends in Cell Biology. 2023;33. 10.1016/j.tcb.2022.04.011.

33. Kriston-Pál É, Czibula Á, Gyuris Z, Balka G, Seregi A, Sükösd F, et al. Characterization and therapeutic application of canine adipose mesenchymal stem cells to treat elbow osteoarthritis. Canadian Journal of Veterinary Research. 2017;81.

34. Pekker E, Priskin K, Szabó-Kriston É, Csányi B, Buzás-Bereczki O, Adorján L, et al. Development of a Large-Scale Pathogen Screening Test for the Biosafety Evaluation of Canine Mesenchymal Stem Cells. Biol Proced Online. 2023;25:33. 10.1186/s12575-023-00226-x.

35. Andrews S, others. FastQC: a quality control tool for high throughput sequence data. 2010. Https://WwwBioinformaticsBabrahamAcUk/Projects/Fastqc/. 2019.

36. Chen S, Zhou Y, Chen Y, Gu J. Fastp: An ultra-fast all-in-one FASTQ preprocessor. In: Bioinformatics. 2018. 10.1093/bioinformatics/bty560.

37. Bray NL, Pimentel H, Melsted P, Pachter L. Near-optimal probabilistic RNA-seq quantification. Nat Biotechnol. 2016;34. 10.1038/nbt.3519.

38. Soneson C, Love MI, Robinson MD. Differential analyses for RNA-seq: transcript-level estimates improve gene-level inferences. F1000Res. 2015;4. 10.12688/f1000research.7563.1.

39. Robinson MD, McCarthy DJ, Smyth GK. edgeR: A Bioconductor package for differential expression analysis of digital gene expression data. Bioinformatics. 2009;26. 10.1093/bioinformatics/btp616.

40. Ritchie ME, Phipson B, Wu D, Hu Y, Law CW, Shi W, et al. Limma powers differential expression analyses for RNA-sequencing and microarray studies. Nucleic Acids Res. 2015;43. 10.1093/nar/gkv007.

41. Wu T, Hu E, Xu S, Chen M, Guo P, Dai Z, et al. clusterProfiler 4.0: A universal enrichment tool for interpreting omics data. Innovation. 2021;2. 10.1016/j.xinn.2021.100141.

42. Zheng GXY, Terry JM, Belgrader P, Ryvkin P, Bent ZW, Wilson R, et al. Massively parallel digital transcriptional profiling of single cells. Nat Commun. 2017;8. 10.1038/ncomms14049.

43. Martin FJ, Amode MR, Aneja A, Austine-Orimoloye O, Azov AG, Barnes I, et al. Ensembl 2023. Nucleic Acids Res. 2023;51:D933–41. 10.1093/nar/gkac958.

44. R Development Core Team. R Core Team (2020). R: A language and environment for statistical computing. R Foundation for Statistical Computing, Vienna, Austria. R Foundation for Statistical Computing. 2019;2.

45. Hao Y, Hao S, Andersen-Nissen E, Mauck WM, Zheng S, Butler A, et al. Integrated analysis of multimodal single-cell data. Cell. 2021;184. 10.1016/j.cell.2021.04.048.

46. Casella G, Munk R, Kim KM, Piao Y, De S, Abdelmohsen K, et al. Transcriptome signature of cellular senescence. Nucleic Acids Res. 2019;47. 10.1093/nar/gkz555.

47. Wang Y, Liu L, Song Y, Yu X, Deng H. Unveiling E2F4, TEAD1 and AP-1 as regulatory transcription factors of the replicative senescence program by multi-omics analysis. Protein Cell. 2022;13. 10.1007/s13238-021-00894-z.

48. Taherian Fard A, Leeson HC, Aguado J, Pietrogrande G, Power D, Gómez-Inclán C, et al. Deconstructing heterogeneity of replicative senescence in human mesenchymal stem cells at single cell resolution. Geroscience. 2024;46. 10.1007/s11357-023-00829-y.

49. Lun ATL, Riesenfeld S, Andrews T, Dao TP, Gomes T, Marioni JC. EmptyDrops: Distinguishing cells from empty droplets in droplet-based single-cell RNA sequencing data. Genome Biol. 2019;20. 10.1186/s13059-019-1662-y.

50. Korsunsky I, Millard N, Fan J, Slowikowski K, Zhang F, Wei K, et al. Fast, sensitive and accurate integration of single-cell data with Harmony. Nat Methods. 2019;16. 10.1038/s41592-019-0619-0.

51. Wickham H. ggplot2: elegant graphics for data analysis. ht tp. had. co. nz/ggplot2/book. Springer; 2009;8.

52. Kevin Blighe, Sharmila Rana, Myles Lewis. EnhancedVolcano: publication-ready volcano plots with enhanced colouring and labeling. 2024.

53. Guo X, Chen L. From G1 to M: a comparative study of methods for identifying cell cycle phases. Briefings in Bioinformatics. 2024;25. 10.1093/bib/bbad517.

54. Tirosh I, Izar B, Prakadan SM, Wadsworth MH, Treacy D, Trombetta JJ, et al. Dissecting the multicellular ecosystem of metastatic melanoma by single-cell RNA-seq. Science (1979). 2016;352. 10.1126/science.aad0501.

55. Wiley CD, Campisi J. The metabolic roots of senescence: mechanisms and opportunities for intervention. Nature Metabolism. 2021;3. 10.1038/s42255-021-00483-8.

56. Deckx S, Heymans S, Papageorgiou AP. The diverse functions of osteoglycin: A deceitful dwarf, or a master regulator of disease. FASEB Journal. 2016;30. 10.1096/fj.201500096R.

57. Freund A, Laberge R-M, Demaria M, Campisi J. Lamin B1 loss is a senescence-associated biomarker. Mol Biol Cell. 2012;23:2066–75. 10.1091/mbc.e11-10-0884.

58. Hang Pham LB, Yoo YR, Park SH, Back SA, Kim SW, Bjørge I, et al. Investigating the effect of fibulin-1 on the differentiation of human nasal inferior turbinate-derived mesenchymal stem cells into osteoblasts. J Biomed Mater Res A. 2017;105. 10.1002/jbm.a.36095.

59. Limbad C, Doi R, McGirr J, Ciotlos S, Perez K, Clayton ZS, et al. Senolysis induced by 25-hydroxycholesterol targets CRYAB in multiple cell types. iScience. 2022;25. 10.1016/j.isci.2022.103848.

60. Zou H, Stoppani E, Volonte D, Galbiati F. Caveolin-1, cellular senescence and age-related diseases. Mech Ageing Dev. 2011;132. 10.1016/j.mad.2011.11.001.

61. Cooley MA, Harikrishnan K, Oppel JA, Miler SF, Barth JL, Haycraft CJ, et al. Fibulin-1 is required for bone formation and Bmp-2-mediated induction of Osterix. Bone. 2014;69:30–8. 10.1016/j.bone.2014.07.038.

62. Zhang D, Liu X, Xu X, Xu J, Yi Z, Shan B, et al. HPCAL1 promotes glioblastoma proliferation via activation of Wnt/β-catenin signalling pathway. J Cell Mol Med. 2019;23. 10.1111/jcmm.14083.

63. Lee BC, Yu KR. Impact of mesenchymal stem cell senescence on inflammaging. BMB Reports. 2020;53. 10.5483/BMBRep.2020.53.2.291.

64. Hao M, Jiang H, Zhao Y, Li C, Jiang J. Identification of potential biomarkers for aging diagnosis of mesenchymal stem cells derived from the aged donors. Stem Cell Res Ther. 2024;15. 10.1186/s13287-024-03689-1.

65. Ruple A, Maclean E, Snyder-Mackler N, Creevy KE, Promislow D. Dog Models of Aging. Annual Review of Animal Biosciences. 2022;10. 10.1146/annurev-animal-051021-080937.

66. Hardwick LJA, Kortum AJ, Constantino-Casas F, Watson PJ. Breed-related expression patterns of Ki67, γH2AX, and p21 during ageing in the canine liver. Vet Res Commun. 2021;45. 10.1007/s11259-020-09784-x.

67. Sándor S, Jónás D, Tátrai K, Czeibert K, Kubinyi E. Poly(A) RNA sequencing reveals age-related differences in the prefrontal cortex of dogs. Geroscience. 2022;44. 10.1007/s11357-022-00533-3.

68. Mirsaidi A, Kleinhans KN, Rimann M, Tiaden AN, Stauber M, Rudolph KL, et al. Telomere length, telomerase activity and osteogenic differentiation are maintained in adipose-derived stromal cells from senile osteoporotic SAMP6 mice. J Tissue Eng Regen Med. 2012;6. 10.1002/term.440.

69. Melk A, Schmidt BMW, Takeuchi O, Sawitzki B, Rayner DC, Halloran PF. Expression of p16INK4a and other cell cycle regulator and senescence associated genes in aging human kidney. Kidney Int. 2004;65. 10.1111/j.1523-1755.2004.00438.x.

70. Zindy F, Quelle DE, Roussel MF, Sherr CJ. Expression of the p16(INK4a) tumor suppressor versus other INK4 family members during mouse development and aging. Oncogene. 1997;15. 10.1038/sj.onc.1201178.

71. Bertolo A, Steffen F, Malonzo-Marty C, Stoyanov J. Canine mesenchymal stem cell potential and the importance of dog breed: Implication for cell-based therapies. Cell Transplant. 2015;24. 10.3727/096368914X685294.

72. A B. Comparative Characterization of Canine and Human Mesenchymal Stem Cells Derived from Bone Marrow. Int J Stem Cell Res Ther. 2015;2. 10.23937/2469-570x/1410005.

73. Li Y, Wu Q, Yujia W, Li L, Bu H, Bao J. Senescence of mesenchymal stem cells (Review). Int J Mol Med. 2017;39. 10.3892/ijmm.2017.2912.

74. Zhou S, Greenberger JS, Epperly MW, Goff JP, Adler C, Leboff MS, et al. Age-related intrinsic changes in human bone-marrow-derived mesenchymal stem cells and their differentiation to osteoblasts. Aging Cell. 2008;7. 10.1111/j.1474-9726.2008.00377.x.

75. Wang D, Xiao F, Feng Z, Li M, Kong L, Huang L, et al. Sunitinib facilitates metastatic breast cancer spreading by inducing endothelial cell senescence. Breast Cancer Research. 2020;22. 10.1186/s13058-020-01346-y.

76. Belcastro E, Rehman AU, Remila L, Park SH, Gong DS, Anton N, et al. Fluorescent nanocarriers targeting VCAM-1 for early detection of senescent endothelial cells. Nanomedicine. 2021;34. 10.1016/j.nano.2021.102379.

77. Alves H, van Ginkel J, Groen N, Hulsman M, Mentink A, Reinders M, et al. A mesenchymal stromal cell gene signature for donor age. PLoS One. 2012;7. 10.1371/journal.pone.0042908.

78. Chelombitko MA, Morgunova G V., Strochkova NY, Zinovkin RA, Pavlyuchenkova AN, Kondratenko ND, et al. Comparative Analysis of Cell Senescence Induced by the Chemotherapeutic Agents Doxorubicin, Cisplatin and Arsenic Trioxide in Human Myoblasts MB135. Advances in Gerontology. 2023;13. 10.1134/S2079057024600010.

79. Li M, Ren H, Zhang Y, Liu N, Fan M, Wang K, et al. MECOM/PRDM3 and PRDM16 Serve as Prognostic-Related Biomarkers and Are Correlated With Immune Cell Infiltration in Lung Adenocarcinoma. Front Oncol. 2022;12. 10.3389/fonc.2022.772686.

80. Bryant AG, Hu M, Carlyle BC, Arnold SE, Frosch MP, Das S, et al. Cerebrovascular Senescence Is Associated With Tau Pathology in Alzheimer’s Disease. Front Neurol. 2020;11. 10.3389/fneur.2020.575953.

81. Qin Z, Chen Z, Weng J, Li S, Rong Z, Zhou C. Elevated HOXA13 expression promotes the proliferation and metastasis of gastric cancer partly via activating Erk1/2. Onco Targets Ther. 2019;12. 10.2147/OTT.S196986.

82. Yu M, Zhan J, Zhang H. HOX family transcription factors: Related signaling pathways and post-translational modifications in cancer. Cellular Signalling. 2020;66. 10.1016/j.cellsig.2019.109469.

83. Lee J, Kim YS, Kim E, Kim Y, Kim Y. Curcumin and hesperetin attenuate d-galactose-induced brain senescence in vitro and in vivo. Nutr Res Pract. 2020;14. 10.4162/nrp.2020.14.5.438.

84. Bérubé NG, Swanson XH, Bertram MJ, Kittle JD, Didenko V, Baskin DS, et al. Cloning and characterization of CRF, a novel C1q-related factor, expressed in areas of the brain involved in motor function. Molecular Brain Research. 1999;63. 10.1016/S0169-328X(98)00278-2.

85. Qiu X, Feng JR, Wang F, Chen PF, Chen XX, Zhou R, et al. Profiles of differentially expressed genes and overexpression of NEBL indicates a positive prognosis in patients with colorectal cancer. Mol Med Rep. 2018;17. 10.3892/mmr.2017.8210.

86. Ni F, Wang F, Li J, Liu Y, Sun X, Chen J, et al. BNC1 deficiency induces mitochondrial dysfunction-triggered spermatogonia apoptosis through the CREB/SIRT1/FOXO3 pathway: the therapeutic potential of nicotinamide riboside and metformin. Biol Reprod. 2024;110. 10.1093/biolre/ioad168.

87. Short SP, Pilat JM, Barrett CW, Reddy VK, Haberman Y, Hendren JR, et al. Colonic Epithelial-Derived Selenoprotein P Is the Source for Antioxidant-Mediated Protection in Colitis-Associated Cancer. Gastroenterology. 2021;160. 10.1053/j.gastro.2020.12.059.

88. Koral K, Paranjpe S, Bowen WC, Mars W, Luo J, Michalopoulos GK. Leukocyte-Specific Protein 1: A novel regulator of hepatocellular proliferation and migration deleted in human hepatocellular carcinoma. Hepatology. 2015;61. 10.1002/hep.27444.

89. Zhang H, Okuyama H, Jain A, Jadhav RR, Wu B, Sturmlechner I, et al. PREX1 improves homeostatic proliferation to maintain a naive CD4+ T cell compartment in older age. JCI Insight. 2024;9. 10.1172/jci.insight.172848.

90. de Resende MF, Vieira S, Chinen LTD, Chiappelli F, da Fonseca FP, Guimarães GC, et al. Prognostication of prostate cancer based on TOP2A protein and gene assessment: TOP2A in prostate cancer. J Transl Med. 2013;11. 10.1186/1479-5876-11-36.

91. Musa J, Aynaud MM, Mirabeau O, Delattre O, Grünewald TGP. MYBL2 (B-Myb): a central regulator of cell proliferation, cell survival and differentiation involved in tumorigenesis. Cell Death and Disease. 2017;8. 10.1038/CDDIS.2017.244.

92. Liu ZM, Bao Y, Li TK, Di Y Bin, Song WJ. MKI67 an potential oncogene of oral squamous cell carcinoma via the high throughput technology. Medicine (United States). 2022;101. 10.1097/MD.0000000000032595.

93. Xi LC, Ji YX, Yin D, Zhao ZX, Huang SC, Yu SL, et al. Effects of Dermatopontin gene silencing on apoptosis and proliferation of osteosarcoma MG-63 cells. Mol Med Rep. 2018;17. 10.3892/mmr.2017.7866.

94. Shandilya J, Roberts SGE. A role of WT1 in cell division and genomic stability. Cell Cycle. 2015;14. 10.1080/15384101.2015.1021525.

95. Tang C, Qiu S, Mou W, Xu J, Wang P. Excessive activation of HOXB13/PIMREG axis promotes hepatocellular carcinoma progression and drug resistance. Biochem Biophys Res Commun. 2022;623. 10.1016/j.bbrc.2022.07.066.

96. Chatsirisupachai K, Palmer D, Ferreira S, de Magalhães JP. A human tissue-specific transcriptomic analysis reveals a complex relationship between aging, cancer, and cellular senescence. Aging Cell. 2019;18. 10.1111/acel.13041.

97. Gruber HE, Hoelscher GL, Ingram JA, Zinchenko N, Hanley EN. Senescent vs. non-senescent cells in the human annulus in vivo: Cell harvest with laser capture microdissection and gene expression studies with microarray analysis. BMC Biotechnol. 2010;10. 10.1186/1472-6750-10-5.

98. Wagner W, Horn P, Castoldi M, Diehlmann A, Bork S, Saffrich R, et al. Replicative senescence of mesenchymal stem cells: A continuous and organized process. PLoS One. 2008;3. 10.1371/journal.pone.0002213.

99. Zou J, Lei T, Guo P, Yu J, Xu Q, Luo Y, et al. Mechanisms shaping the role of ERK1/2 in cellular senescence (Review). Mol Med Rep. 2019;19. 10.3892/mmr.2018.9712.

100. Fujita M, Sasada M, Iyoda T, Fukai F. Involvement of Matricellular Proteins in Cellular Senescence: Potential Therapeutic Targets for Age-Related Diseases. International Journal of Molecular Sciences. 2024;25. 10.3390/ijms25126591.

101. Yeo EJ. Hypoxia and aging. Experimental and Molecular Medicine. 2019;51. 10.1038/s12276-019-0233-3.

102. Xu W, Liu X, Han W, Wu K, Zhao M, Mei T, et al. Inhibiting HIF-1 signaling alleviates HTRA1-induced RPE senescence in retinal degeneration. Cell Communication and Signaling. 2023;21. 10.1186/s12964-023-01138-9.

103. Sun Y, Yu X, Gao X, Zhang C, Sun H, Xu K, et al. RNA sequencing profiles reveal dynamic signaling and glucose metabolic features during bone marrow mesenchymal stem cell senescence. Cell Biosci. 2022;12. 10.1186/s13578-022-00796-5.

104. Nassrally MS, Lau A, Wise K, John N, Kotecha S, Lee KL, et al. Cell cycle arrest in replicative senescence is not an immediate consequence of telomere dysfunction. Mech Ageing Dev. 2019;179. 10.1016/j.mad.2019.01.009.

105. Perez K, Ciotlos S, McGirr J, Limbad C, Doi R, Nederveen JP, et al. Single nuclei profiling identifies cell specific markers of skeletal muscle aging, frailty, and senescence. Aging. 2022;14. 10.18632/aging.204435.

106. Moussavi-Harami F, Duwayri Y, Martin JA, Moussavi-Harami F, Buckwalter JA. Oxygen effects on senescence in chondrocytes and mesenchymal stem cells: consequences for tissue engineering. Iowa Orthop J. 2004;24.

107. Kawamura H, Nakatsuka R, Matsuoka Y, Sumide K, Fujioka T, Asano H, et al. TGF-β Signaling Accelerates Senescence of Human Bone-Derived CD271 and SSEA-4 Double-Positive Mesenchymal Stromal Cells. Stem Cell Reports. 2018;10. 10.1016/j.stemcr.2018.01.030.

108. Volonte D, Galbiati F. Caveolin-1, a master regulator of cellular senescence. Cancer and Metastasis Reviews. 2020;39. 10.1007/s10555-020-09875-w.

109. Fittipaldi S, Mercatelli N, Dimauro I, Jackson MJ, Paronetto MP, Caporossi D. Alpha B-crystallin induction in skeletal muscle cells under redox imbalance is mediated by a JNK-dependent regulatory mechanism. Free Radic Biol Med. 2015;86. 10.1016/j.freeradbiomed.2015.05.035.

110. Masamha CP, Benbrook DM. Cyclin D1 degradation is sufficient to induce G1 cell cycle arrest despite constitutive expression of cyclin E2 in ovarian cancer cells. Cancer Res. 2009;69. 10.1158/0008-5472.CAN-09-0913.

111. Leontieva O V., Demidenko ZN, Blagosklonny M V. MEK drives cyclin D1 hyperelevation during geroconversion. Cell Death Differ. 2013;20. 10.1038/cdd.2013.86.

112. Pagano M, Theodoras AM, Tam SW, Draetta GF. Cyclin D1-mediated inhibition of repair and replicative DNA synthesis in human fibroblasts. Genes Dev. 1994;8. 10.1101/gad.8.14.1627.

113. Duangprom S, Kheolamai P, Tantrawatpan C, Manochantr S. High glucose inhibits proliferation, migration, and osteogenic differentiation of human placenta-derived mesenchymal stem cells. Sci Rep. 2025;15. 10.1038/s41598-025-06454-3.

114. Laphanuwat P, Likasitwatanakul P, Sittithumcharee G, Thaphaengphan A, Chomanee N, Suppramote O, et al. Cyclin D1 depletion interferes with oxidative balance and promotes cancer cell senescence. J Cell Sci. 2018;131. 10.1242/jcs214726.

115. Jazbutyte V, Fiedler J, Kneitz S, Galuppo P, Just A, Holzmann A, et al. MicroRNA-22 increases senescence and activates cardiac fibroblasts in the aging heart. Age (Omaha). 2013;35. 10.1007/s11357-012-9407-9.

116. Chen X, Chen J, Xu D, Zhao S, Song H, Peng Y. Effects of Osteoglycin (OGN) on treating senile osteoporosis by regulating MSCs. BMC Musculoskelet Disord. 2017;18. 10.1186/s12891-017-1779-7.

